# ATG8ylation of vacuolar membrane protects plants against cell wall damage

**DOI:** 10.1101/2024.04.21.590262

**Authors:** Jose Julian, Peng Gao, Alessia Del Chiaro, Juan Carlos De La Concepcion, Laia Armengot, Marc Somssich, Heloise Duverge, Marion Clavel, Nenad Grujic, Roksolana Kobylinska, Ingo Polivka, Maarten Besten, Christian Dank, Barbara Korbei, Andreas Bachmair, Nuria S. Coll, Joris Sprakel, Yasin Dagdas

## Abstract

Vacuoles are essential for cellular metabolism, growth, and the maintenance of internal turgor pressure. They sequester lytic enzymes, ions, and secondary metabolites that, if leaked into the cytosol, could lead to cell death. Despite their pivotal roles, quality control pathways that safeguard vacuolar integrity remained elusive in plants. Here, we discovered a conserved vacuolar quality control (VQC) pathway that is activated upon cell wall damage in a turgor pressure dependent manner. Cell wall perturbations induce a distinct modification – ATG8ylation – on the vacuolar membrane (tonoplast) that is regulated by the V-ATPase and ATG8 conjugation machinery. Genetic disruption of tonoplast ATG8ylation impairs vacuolar integrity, leading to cell death. Together, our findings reveal a homeostatic pathway that preserves vacuolar integrity upon cell wall damage.

## Introduction

Plant cells are pressurized compartments, encased within a rigid wall that must simultaneously resist turgor pressure and accommodate growth through flexibility (*1, 2*). The cell wall, a dynamic matrix of structural polysaccharides including cellulose, pectin, and lignin, along with a suite of structural and modulatory proteins, is critical for defining cell morphology and maintaining cellular integrity (*3, 4*). Moreover, it serves as the primary defense barrier against pathogens and abiotic stresses (*4–6*). Consistent with its role in cellular viability and function, plants have evolved elaborate cell wall integrity (CWI) pathways that closely surveil disruptions in wall composition and mechanics to initiate homeostatic mechanisms that restore cell wall integrity. (*2, 6*). So far, CWI studies focused on cell wall homeostasis. The impact of cell wall damage on cellular organelles and the contribution of these organelles to maintaining cell wall integrity remain elusive. (*4*).

Vacuoles account for up to 80% of the cellular volume (*7, 8*). The expansion of the vacuole within the inflexible cell wall generates turgor pressure that is characteristic of plant cells and required for growth (*9–11*). The receptor-like kinase FERONIA, a key sensor of cell wall integrity, has been implicated in the modulation of vacuolar morphology during cell expansion (*9*), suggesting a potential link between the cell wall integrity sensing and vacuolar integrity. Furthermore, cell wall integrity signaling is sensitive to osmotic fluctuations; for instance, osmolytes like sorbitol can dampen CWI responses (*12*). A pressing question that remains unexplored is how plants preserve vacuolar integrity when faced with sudden changes in cell wall structure. Without the protection of the rigid cell wall, the turgor pressure imbalance could lead to the rupture of the tonoplast and cell death (*13*). Therefore, elucidating quality control pathways that safeguard vacuolar integrity is essential to improve plant resilience and adaptability.

A hallmark of macroautophagy (hereafter autophagy) is the conjugation of the ubiquitin-like protein ATG8 to the double-membraned phagophore via the concerted action of conserved Autophagy Related Gene (ATG) proteins (*14*). While investigating how cell wall damage influence autophagy, unexpectedly, we discovered that cell wall damage triggers conjugation of ATG8 to the single membraned tonoplast in both *Arabidopsis thaliana* and *Marchantia polymorpha*. Genetic characterization of this Conjugation of ATG8 to Single Membrane (CASM) process revealed V-ATPase-ATG16 axis as a key module regulating tonoplast ATG8ylation. Ultrastructural analysis of CASM deficient plants have shown that tonoplast ATG8ylation is essential for vacuolar integrity and cell viability. Altogether, our findings reveal ATG8ylation is a vacuolar quality control (VQC) mechanism that protects vacuolar integrity upon cell wall damage.

## Results

### Cell wall damage triggers ATG8ylation of the tonoplast

To investigate if cell wall damage triggers autophagy, we performed confocal microscopy analysis on ATG8 reporter lines upon cell wall damage. First, we tested the co-localization of mCherry-ATG8A with the tonoplast markers VAMP711-YFP and γ-TIP-GFP under control conditions (Fig. 1A and fig. S1A). Few autophagic puncta that we detected did not colocalize with the tonoplast under basal conditions (R^PEARSON^=0.12±0.05, R^SPEARMAN^=0.15±0.07) (Fig. 1A and fig. S1A).

**Figure 1.**
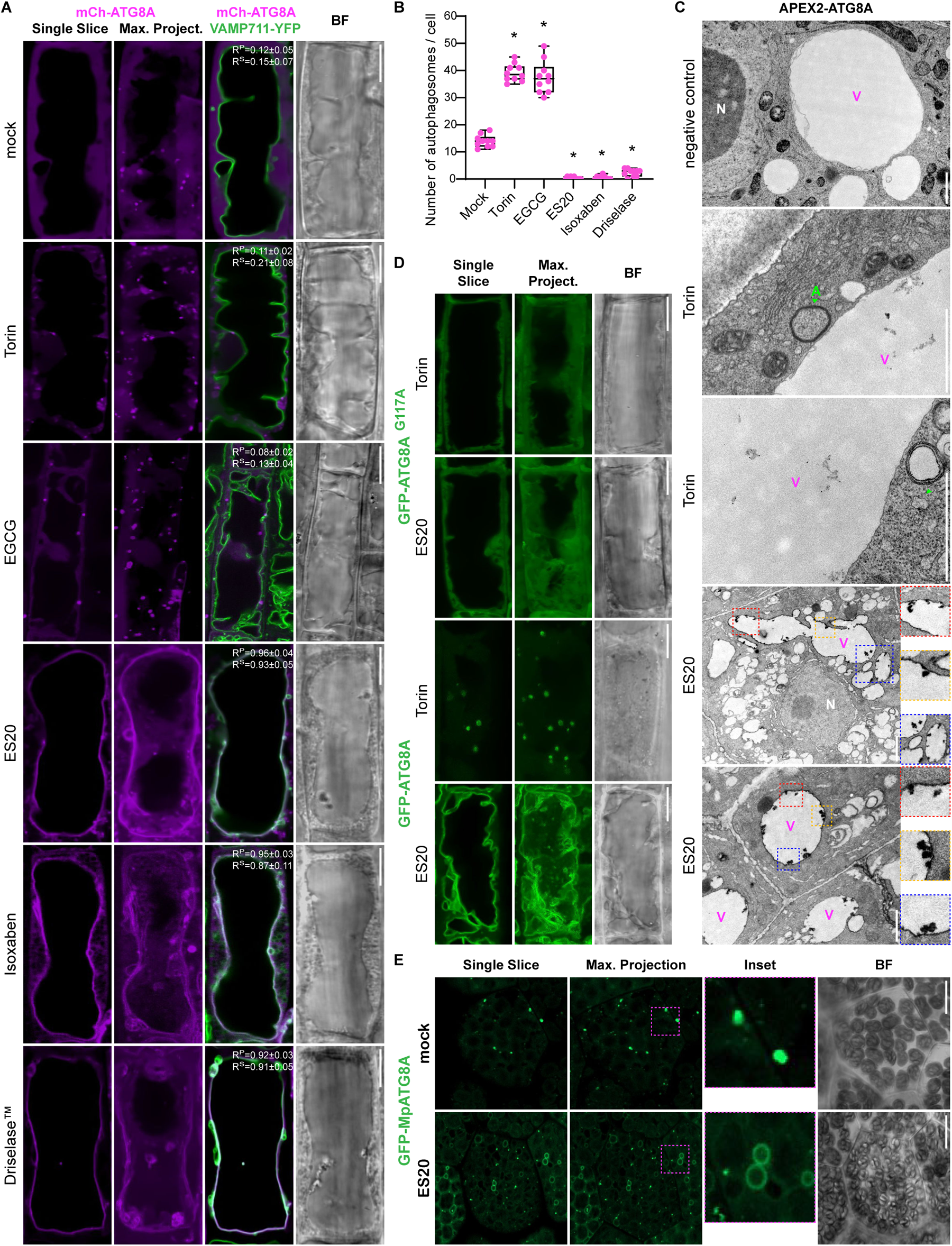
Cell wall damage induces ATG8ylation of the tonoplast. **(A)** Confocal micrographs of root cells in the early elongation zone of *Arabidopsis thaliana*, highlighting the localization of mCherry-ATG8A (in magenta) to illustrate re-localization of ATG8 to the tonoplast upon cell wall damage. The panel includes a single optical slice and a maximum intensity projection of a whole cell (20 µm depth), alongside a merged image with VAMP711-YFP (tonoplast marker) and a corresponding bright field image. Scale bar, 10 µm. Pearson and Spearman co-localization values are presented, showcasing the association between ATG8A and the tonoplast. Treatment conditions include mock, Torin (1.5 hours, 9 µM), EGCG (30 minutes, 50 µM), ES20 (8 hours, 100 µM), Isoxaben (3 days, 3 nM), and Driselase (1 hour, 1%). **(B)** Quantification of autophagosomes under treatment conditions depicted in Fig. 1A. Statistical analysis was performed using Wilcoxon test to compare each treatment condition against the mock, indicating significant differences at p-values lower than 0.01. **(C)** Electron microscopy (EM) images displaying APEX2-ATG8A localization post-Torin (1.5 hours, 9 µM) or ES20 treatments (8 hours, 100 µM), with or without DAB staining (negative control) to highlight APEX2-ATG8A signal. Torin treated samples show typical autophagosome structures, whereas ES20 treatments lead to the labeling of the tonoplast. Insets show densely labeled tonoplast invaginations. N: nucleus, V and magenta arrowheads: vacuole, A and green arrowheads: autophagosome. Scale bar, 1 µm. **(D)** Confocal micrographs of *Arabidopsis thaliana* root cells expressing the GFP-ATG8A^G117A^ mutant, which is incapable of conjugating to membranes. Images are shown after treatment with Torin (1.5 hours, 9 µM) or ES20 (8 hours, 100 µM), following the same pattern of a single optical slice, a maximum intensity projection, and a bright field image. Scale bar, 10 µm. **(E)** Confocal micrographs of *Marchantia polymorpha*, comparing GFP-ATG8A localization under mock and ES20 (8 hours, 100 µM) treatment conditions. Scale bar, 10 µm.

We then used TOR kinase inhibitor Torin1 (hereafter Torin) to induce bulk autophagy (*15*). Torin increased the number of the autophagic puncta, but these puncta also did not colocalize with the tonoplast (R^P^=0.11±0.02, R^S^=0.21±0.08) (Fig. 1, A and B, and fig. S1). Treatment with the pectin methylesterase inhibitor epigallocatechin gallate (EGCG), which increases cell wall stiffness by changing pectin methylation and polymerization (*9*), also increased the number of autophagic puncta without any significant colocalization with the tonoplast (R^P^=0.08±0.02, R^S^=0.13±0.04) (Fig. 1, A and B, and fig. S1).

In contrast, when we inhibit cellulose biosynthesis and loosen the cell wall with ES20 or Isoxaben treatments (*16*), we observed loss of autophagic puncta and a notable re-localization of mCherry-ATG8A to the tonoplast (ES20: R^P^=0.96±0.04, R^S^=0.93±0.05; Isoxaben: R^P^=0.95±0.03, R^S^=0.87±0.11). Similarly, treatment with Driselase™, a fungal enzymatic cocktail that is used to mimic fungal infection (*12*), also led to the labeling of tonoplast with mCherry-ATG8A (R^P^=0.92±0.03, R^S^=0.91±0.05) (Fig. 1, A and B, and fig. S1). Tonoplast labeling upon cell wall damage is not specific to the ATG8A isoform, since all nine Arabidopsis GFP-labeled ATG8 isoforms showed similar tonoplast re-localization upon ES20 treatment (Fig. S2A).

Next, we implemented Ascorbate Peroxidase 2 (APEX2)-based electron microscopy to study tonoplast ATG8ylation at the ultrastructural level. APEX2 is a plant peroxidase enzyme that has been engineered as a genetically encoded electron microscopy tag. Addition of the substrate Diaminobenzidine (DAB) leads to localized ROS production and increased contrast in electron micrographs (*17*). Without DAB, there was minimal background labeling (Fig. 1C). Addition of the substrate led to some background labeling at the Golgi apparatus, suggesting inherent plant peroxidase activity could lead to the labeling of the Golgi apparatus (fig. S3). Nevertheless, the significant increase in the contrast prompted us to test the APEX2-ATG8A samples upon Torin and ES20 treatments. We could visualize typical autophagosome structures in Torin treated samples (Fig. 1C). In contrast, upon ES20 treatment, APEX2 labeled the tonoplast (Fig. 1C). In micrographs obtained from ES20 treated samples, we could also detect invaginations of the tonoplast that are densely labeled with APEX2-ATG8A (Fig. 1C). Altogether, consistent with our confocal microscopy results, APEX2 based electron microscopy showed re-localization of ATG8A to the tonoplast upon cell wall damage.

During autophagosome biogenesis, ATG8 is conjugated to the growing double membraned phagophore via a lipid modification. Through a ubiquitination-like reaction, the C-terminal glycine residue gets lipidated for membrane insertion (*18*). To test if ATG8 is conjugated to the tonoplast, we tested relocation of mutant ATG8A^G117A^ that cannot get lipidated. This mutant had a diffuse pattern and failed to relocate to the tonoplast upon ES20 treatment (Fig. 1D). Collectively, these findings suggest that cell wall damage induces CASM at the tonoplast.

Finally, we investigated the evolutionary conservation of this pathway by testing if cell wall damage induces tonoplast ATG8ylation in the early diverging land plant *Marchantia polymorpha*. We labeled Marchantia GFP-MpATG8A and GFP-MpATG8B and showed that upon ES20 treatment both ATG8 isoforms relocate from puncta-like autophagosomes to the tonoplast (Fig. 1E and fig. S2B). These results suggest cell wall damage-induced tonoplast ATG8ylation is conserved across land plants.

### Genetic basis of tonoplast ATG8ylation

Next, we sought to dissect the genetic basis of tonoplast ATG8ylation. Previous studies in mammalian cells have shown that unlike autophagy, CASM requires only a subset of the ATG proteins (*19*). We generated stable transgenic Arabidopsis lines that express GFP-ATG8A in the mutant background of the core ATG protein ATG5, ATG11, and ATG16. As expected, autophagosome formation is blocked in all three mutant backgrounds (Fig. 2A). In contrast, tonoplast ATG8ylation is only inhibited in *atg5* and *atg16* mutants, which are part of the ATG8 conjugation machinery. While in *atg11* mutant, which is part of the ATG1 kinase complex, tonoplast ATG8ylation is not affected (Fig. 2A). This suggests tonoplast ATG8ylation requires the ATG8 conjugation machinery but is independent of the ATG1 kinase complex.

**Figure 2.**
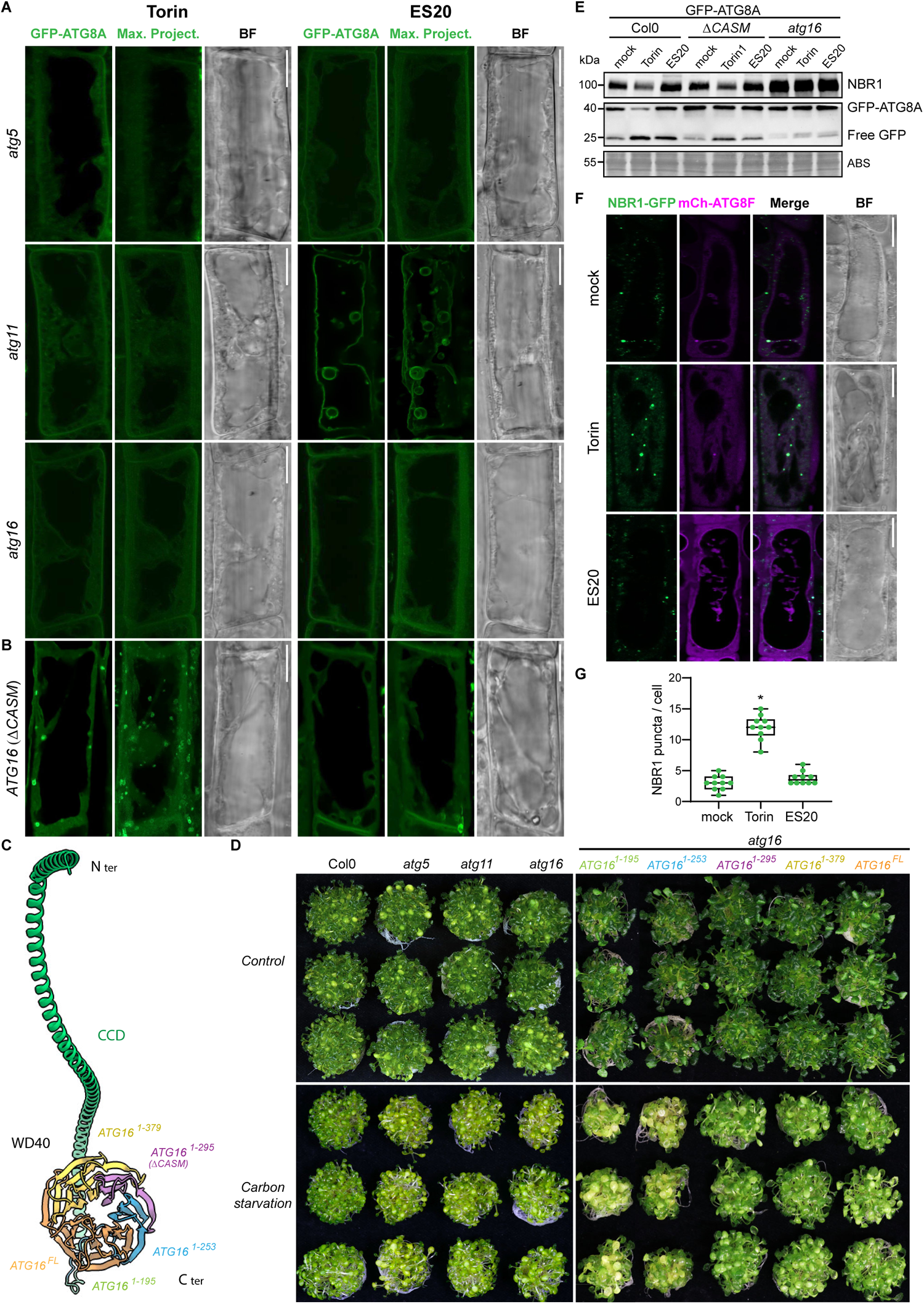
Genetic basis of tonoplast ATG8ylation. **(A)** Confocal micrographs of GFP-ATG8A expressed in *atg5*, *atg11*, and *atg16* mutant backgrounds of *Arabidopsis thaliana* root cells treated with Torin (1.5 hours, 9 µM) or ES20 (8 hours, 100 µM). Each panel includes a single optical slice, a maximum intensity projection of the entire cell (20 µm depth), and a corresponding bright field image to illustrate ATG8A behavior in various genetic backgrounds. Scale bar, 10 µm. **(B)** Similar confocal imaging setup as in Fig 2A, focusing on the *atg16* mutant background complemented with an ATG16 variant (*ΔCASM*), which retains canonical autophagy functionality but lacks the capacity for non-canonical autophagy due to the absence of conjugation of ATG8 to single membranes (CASM). Scale bar, 10 µm. **(C)** Schematic representation of ATG16 protein structure highlighting its critical domains: CCD and WD40. The figure details various truncations of ATG16 used to complement the *atg16* mutant, represented in different colors to indicate the portions of the protein retained (1-195 in green, 1-253 in blue, 1-295 in purple—designated *ΔCASM*, 1-379 in yellow, and the full-length protein in orange). **(D)** Carbon starvation assay results juxtaposed with normal growth conditions across three replicates for wild-type (Col0), *atg5*, *atg11*, *atg16* mutants, and complemented *atg16* lines with various ATG16 truncations. *ΔCASM* line and longer complemented variants show no sensitivity to carbon starvation, unlike the *atg16* mutant or other *atg* mutants. **(E)** Western blot analysis of plant material expressing GFP-ATG8A in Col0, *Δ CASM/atg16*, and *atg16* backgrounds under mock, Torin (4 hours, 9 µM), or ES20 (8 hours, 100 µM) treatments. Western blots are probed with anti-NBR1 and anti-GFP. Amido black staining used as for loading control. Statistical analysis was performed using Wilcoxon test to compare each treatment condition against the mock, indicating significant differences at p-values lower than 0.01. **(F)** Confocal micrographs showing NBR1-GFP localization (in green) and mCh-ATG8F (in magenta) in Arabidopsis root cells, under mock, Torin (1.5 hours, 9 µM), and ES20 (8 hours, 100 µM) treatments. Images include separate channels for NBR1-GFP and mCh-ATG8F, a merged image, and a corresponding bright field image, providing insight into the colocalization and dynamics of autophagy markers under stress conditions. Scale bar, 10 µm. **(G)** Quantification of NBR1 puncta across treatments. Statistically significant differences compared to mock treatment are marked with an asterisk, denoting a p-value of 0.01.

ATG16 plays a key role in determining the site of ATG8 conjugation. During autophagy, ATG16 coiled-coil domain (CCD) interact with phosphatidylinositol 3-phosphate (PI3P) binding protein WIPI2 to conjugate ATG8 to the growing phagophore (*20, 21*). During CASM, the WD40 domain mediates ATG8 conjugation to single membranes (*22*). Given ATG16’s involvement in spatial distribution of ATG8 conjugation, we hypothesized that complementation of *atg16* mutant with truncated ATG16 variants could provide a genetic tool where autophagy could still happen, but CASM is inhibited. To test this, we complemented the *atg16* mutant with different C-terminal truncations of ATG16 where the WD40 domain is located, and tested autophagic flux, carbon starvation sensitivity, and autophagosome biogenesis (Fig. 2, B to E). Complementation with ATG16 1-295 (hereafter *ΔCASM*), restored canonical autophagy: *(i)* GFP-ATG8A formed punctate structures upon Torin treatment (Fig. 2B), *(ii)* unlike the *atg16* or *atg5* mutants, complemented seedlings were insensitive to carbon starvation (Fig. 2D), and *(iii)* there was an increase in free GFP and a decrease in the autophagy receptor NBR1 protein (*23*) levels upon Torin treatment, indicative of functional autophagic flux (Fig. 2E). In contrast, tonoplast ATG8ylation is inhibited in *ΔCASM* cells (Fig. 2B). These results suggest ATG16 WD40 domain is essential for CASM in Arabidopsis. Crucially, the *ΔCASM* line provides us a genetic tool that bypasses the pleiotropic effects seen with *atg* mutants, thus allowing for targeted investigation into the physiological and cellular significance of tonoplast ATG8ylation following cell wall damage.

Next, we tested whether all autophagic processes are re-routed to the tonoplast, by checking the localizations of the archetypical selective autophagy receptor NBR1 (*23*) and the recently characterized plant selective autophagy adaptor, CFS1 (*24*). Both proteins interact with ATG8 directly via their conserved ATG8 interacting motifs (*23, 24*). NBR1 is located within the autophagosomes together with its cargo. Whereas CFS1 is located on the outer autophagosome membrane and interacts with ESCRT machinery for autophagosome sorting (*23, 24*). Although both proteins responded to Torin treatment and formed more punctate structures, they did not relocate to the tonoplast upon ES20 treatment (Fig. 2, F and G, and fig. S4). Notably, although rather weak, we observed some CFS1 vacuolar localization upon ES20 treatment (fig. S4A). Since CFS1 is located on the outer autophagosome membrane, blockage of the autophagic flux could lead to weak vacuolar localization of this protein. Nevertheless, these findings indicate cell wall damage does not re-route all autophagic processes, rather triggers selective ATG8ylation of the tonoplast.

Since the cell wall integrity sensor FERONIA and its interacting partner Leucine Rich Repeat Extensin (LRX) family were previously shown to link cell wall integrity to vacuolar morphology (*9*), we next tested if FERONIA regulates tonoplast ATG8ylation. We expressed GFP-ATG8A in *fer-4* and *lrx3/4/5* mutants. GFP-ATG8A localization after Torin or ES20 treatment was indistinguishable from the wild-type Col0 background (fig. S5), indicating that FERONIA signaling pathway does not function in tonoplast ATG8ylation.

Together, these findings demonstrate that CASM and canonical autophagy are two independent pathways that are regulated by different protein networks.

### What is the role of tonoplast ATG8ylation?

Our main hypothesis is that cell wall damage will weaken the counterbalance that the cell wall provides against the turgor pressure contained within the vacuole and threaten vacuolar integrity. First, we tested this hypothesis by combining ES20 treatments with osmolyte sorbitol. Consistent with our hypothesis, Sorbitol treatment suppressed tonoplast ATG8ylation upon ES20 treatment (Fig. 3A).

**Figure 3.**
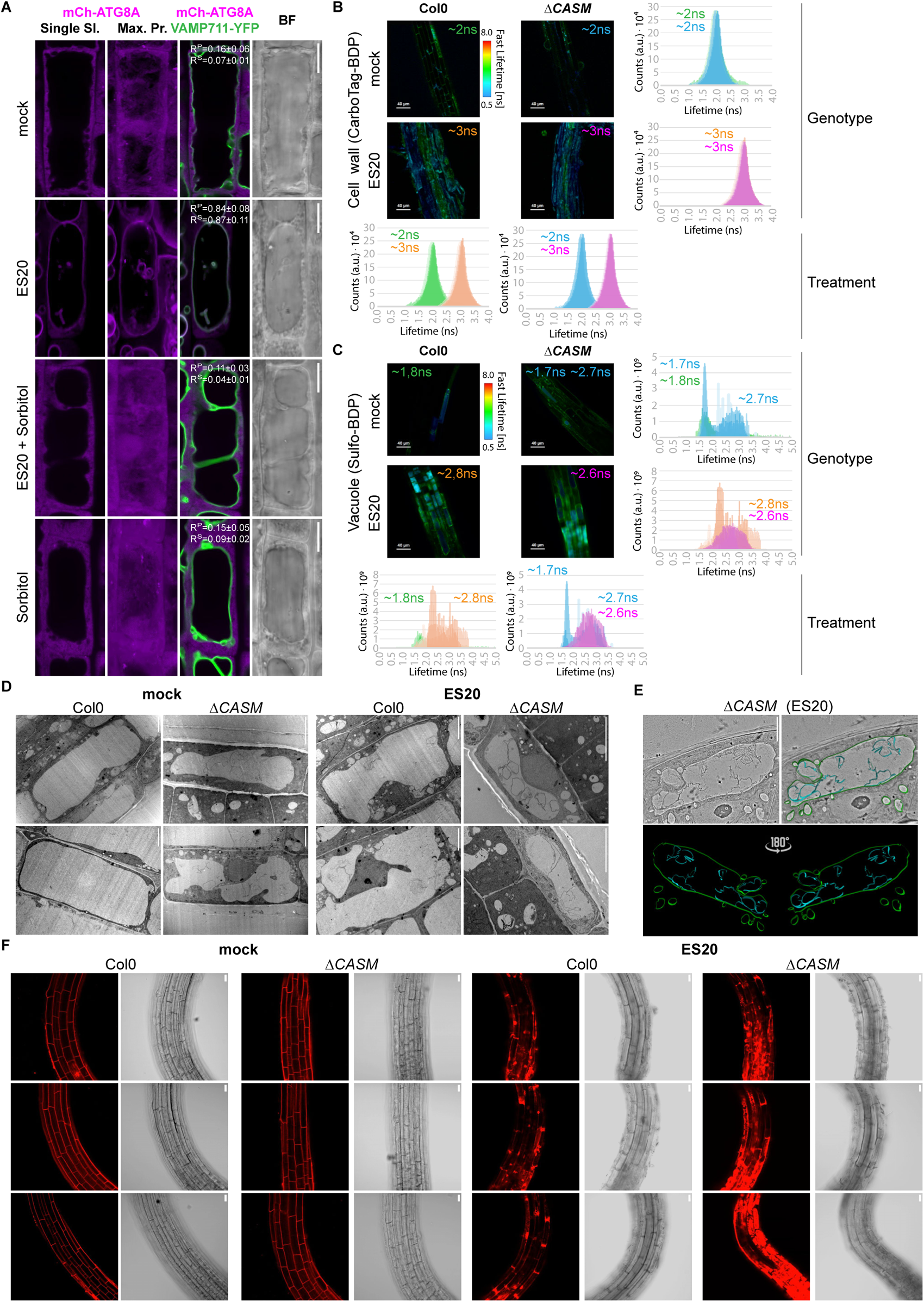
Tonoplast ATG8ylation maintains vacuolar integrity upon cell wall damage in a turgor pressure-dependent manner. **(A)** Confocal micrographs of Arabidopsis root cells expressing mCh-ATG8A and VAMP711-YFP, highlighting ATG8A localization and tonoplast integrity under mock, ES20 (8 hours, 100 µM), sorbitol (8 hours, 50 mM), and combined sorbitol (8 hours, 50 mM) + ES20 (8 hours, 100 µM) treatments. The sequence includes mCh-ATG8A in a single slice, a maximum intensity projection, a merged image of mCh-ATG8A and VAMP711-YFP channels, and a corresponding bright field image. Pearson and Spearman co-localization analyses are presented to quantify co-localization under each condition. Scale bar, 10 µm. **(B)** Fluorescence lifetime imaging microscopy (FLIM) analysis of Col0 and *ΔCASM* backgrounds treated with CarboTag-BDP, a fluorescent cell wall mechanoprobe, for 30 minutes at 10 µM concentration, following mock or ES20 (8 hours, 100 µM) treatments. The fluorescence lifetime of the probe across three biological replicates, with average lifetimes reported in nanoseconds. Four comparative graphs detail the lifetime variance per treatment and genotype. Scale bar, 40 µm. **(C)** FLIM analysis utilizing Sulfo-BDP, a vacuolar mechanoprobe, to assess vacuolar crowding under the same conditions. Lifetime measurements in nanoseconds highlight differences in vacuolar crowding between treatments and genetic backgrounds. Average Lifetime was measured for all values below 2ns (Peak 1), and above 2 ns (Peak 2). Scale bar, 40 µm. **(D)** Transmission electron micrographs demonstrating vacuolar morphology changes in Col0 and *ΔCASM* backgrounds under mock and ES20 (8 hours, 100 µM) treatments. The images reveal the significant fragmentation and invaginations upon cell wall damage in *ΔCASM* line. Scale bar, 5 µm. **(E)** Electron Tomography analysis of *ΔCASM* root cell treated with ES20 (8 hours, 100 µM), providing a detailed 3-dimensional visualization of vacuolar morphology and the surrounding cellular environment. The tomogram is presented with a 180° rotation to enhance structural observation. Scale bar, 5 µm. **(F)** Propidium Iodide (PI) staining of root cells from Col0 and *ΔCASM* backgrounds under mock and ES20 (8 hours, 100 µM) treatments, assessing cell viability and membrane integrity. Three replicates are shown for each genotype and treatment. Scale bar, 10 µm.

Next, we sought to determine whether cell wall damage affects cellular mechanics by taking advantage of the recently established cell wall, vacuole, and cytoplasm targeted BODIPY-based mechanoprobes (*25*). First, we measured the fluorescence lifetime of the cell wall porosity reporter CarboTag-BDP upon ES20 treatment in the wild-type Col0 and *ΔCASM* lines. CarboTag-BDP probe had increased lifetime in both Col0 and *ΔCASM* seedlings (mock ∼2 ns, ES20 ∼3 ns), indicative of decreased cell wall porosity, demonstrating cell wall defects caused by ES20 (Fig. 3B). We then tested the fluorescence lifetime of the vacuolar mechanoprobe Sulfo-BDP to test molecular crowding inside of the vacuole. Excitingly, Sulfo-BDP showed 2 peaks in our lifetime measurements (first peak below 2 ns and second peak after 2 ns). *ΔCASM* seedlings already had vacuoles emitting higher lifetime signals under control conditions (first peak: Col0 ∼1.7 ns, *ΔCASM* ∼1.8 ns; second peak: Col0 none, *ΔCASM* ∼2.6 ns), suggesting the vacuoles are already damaged in *ΔCASM* cells (Fig. 3C). Upon ES20 treatment all vacuoles had increased lifetimes in both genotypes tested (second peak: Col0 ∼2.6 ns, *ΔCASM* ∼2.6 ns) further demonstrating the vacuolar damage triggered by cell wall damage (Fig. 3C). Finally, we tested the cytoplasmic mechanoprobe PEG-BDP upon ES20 treatment. Both Col0 and *ΔCASM* had increased lifetimes (mock: Col0 ∼3.5 ns, *ΔCASM* ∼3.7 ns; ES20: Col0 ∼3.8 ns, *ΔCASM* ∼4.1 ns) (fig. S6). However, PEG-BDP fluorescence lifetime was consistently higher in *ΔCASM* compared to Col0, suggesting cytosolic viscosity is more severely affected in cells that lack tonoplast ATG8ylation, which is indicative of defects in vacuolar maintenance of turgor mechanostasis (fig. S6). In depth studies are necessary to understand the molecular basis of the increased cytosolic viscosity in the *ΔCASM* line, but these findings prompted us to further investigate vacuolar integrity upon cell wall damage.

To further assess vacuolar integrity, we visualized vacuolar morphology using transmission electron microscopy and electron tomography. Compared to the mock condition, ES20 treatment induced vacuolar invaginations in the wild-type Col0 background (Fig. 3D). The *ΔCASM* mutant already showed vacuolar fragmentation and invaginations under control conditions, which became more severe upon ES20 treatment (Fig. 3, D and E, and movie S1). This led in some occasions to an extreme phenotype featuring vacuoles traversing to the adjacent cells through the holes formed upon ES20 treatment (fig. S7).

Finally, we tested the contribution of tonoplast ATG8ylation to cellular integrity with Propidium Iodide (PI) staining. Propidium iodide is impermeable to living cells and commonly used as a cell viability assay (*12, 26*). Upon ES20 treatment, we observed an increase in PI staining, indicative of increased cell death, in both Col0 and *ΔCASM* mutant (Fig. 3F and fig. S8). However, cell death after ES20 treatment was significantly exacerbated in the *ΔCASM* mutant. Altogether, these findings demonstrate tonoplast ATG8ylation is essential to maintain vacuolar integrity and cell survival upon cell wall damage.

### Molecular basis of tonoplast ATG8ylation

Membrane repair is typically coordinated by the ESCRT complex (*27–30*). To investigate a possible link between ESCRT and cell wall damage-induced tonoplast ATG8ylation, we visualized key ESCRT proteins FREE1, ALIX, and VPS23 upon ES20 treatment (*31*). We observed no significant alterations in the localization of these proteins (fig. S9), suggesting that the ESCRT machinery is not involved in tonoplast ATG8ylation triggered by cell wall damage.

But how does cell wall damage induce conjugation of ATG8 to the tonoplast? Since the acidic pH is central to vacuolar function, we hypothesized perturbations in vacuolar pH could signal for tonoplast ATG8ylation. To test this, we measured the vacuolar pH upon ES20 treatment using the LysoSensor™ probe, which only shows fluorescence signal at 440 nm (blue) in compromised vacuoles (*32*). ES20 treatment increased the vacuolar pH (Fig. 4A), suggesting vacuolar pH could be the link between cell wall damage and tonoplast ATG8ylation. To directly test if the increase in vacuolar pH is the driver of tonoplast ATG8ylation, we used the ionophore monensin that serves as a proton-sodium antiporter and increases vacuolar pH (*33*). Monensin mimicked cell wall damage and *(i)* induced tonoplast ATG8ylation (fig. S10A), *(ii)* reduced autophagic puncta (fig. S10B), *(iii)* and blocked autophagic flux of NBR1 (fig. S10C). Monensin-induced tonoplast ATG8ylation is also conserved in Marchantia (fig. S10D). Monensin induced conjugation of all nine Arabidopsis ATG8 isoforms to the tonoplast (fig. S11A). Finally, we showed that ATG8 conjugation is also essential for monensin-induced tonoplast ATG8ylation (fig. S11B). A time course analysis of GFP-ATG8A upon monensin treatment revealed that within ∼20 mins, ATG8 is conjugated to the tonoplast, which is followed by fragmentation of the tonoplast in ∼60 mins (fig. S11C). This aligns with recent findings that demonstrate the ability of ATG8 to alter the morphology of the membranes it attaches to (*34*), indicating that ATG8-mediated restructuring of the tonoplast is vital for preserving vacuolar integrity under stress conditions.

**Figure 4.**
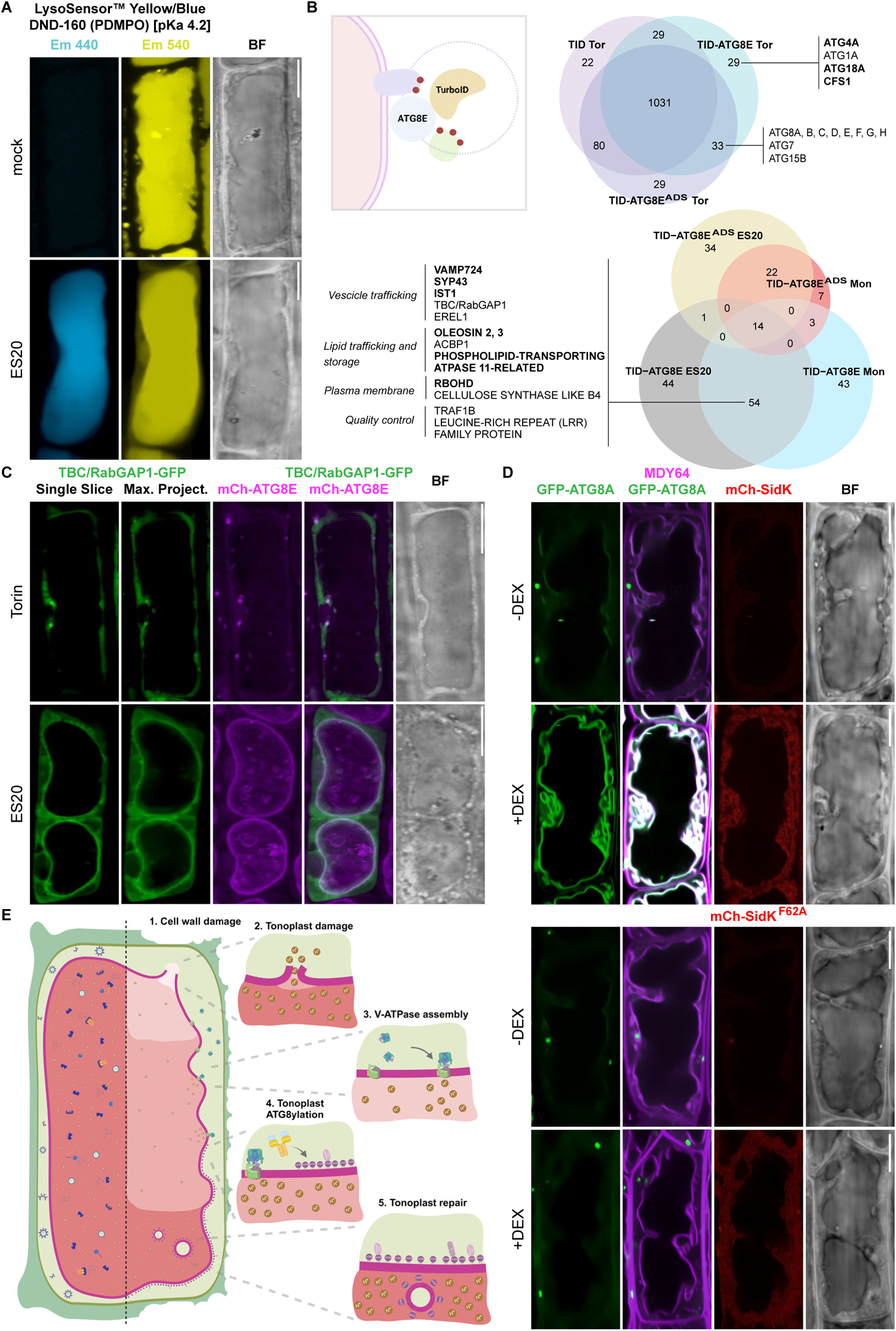
Molecular basis of tonoplast ATG8ylation. **(A)** Cell wall damage increases vacuolar pH. Confocal images of Col0 roots treated with LysoSensor™ Yellow/Blue DND-160 under mock or ES20 (8 hours, 100 µM) conditions, showcasing the probe’s dual emission in yellow and blue, which vary in intensity based on pH. Scale bar, 10 µm. **(B)** ATG8a proxitome upon cell wall damage and CASM activation. Venn diagram summarizing the results of a TurboID-based proximity-labeling proteomics experiment under Torin (1.5 hours, 9 µM), Monensin (0.5 hours, 200 µM), and ES20 (8 hours, 100 µM) treatments, highlighting the overlap and unique proteins identified across conditions. Bold letters indicate the protein is exclusively enriched under that condition. ATG8A-ADS mutant and TurboID alone are used as negative controls. **(C)** Confocal micrographs showing the localization of TBC/RabGAP1-GFP and mCh-ATG8E in *Arabidopsis thaliana* root cells, under mock or ES20 (8 hours, 100 µM) treatments. Each set comprises a single optical slice and a maximum intensity projection for both proteins, a merge image of the maximum projections and a corresponding bright field image. Scale bar, 10 µm. **(D)** SidK expression induces tonoplast ATG8ylation. Confocal micrographs of GFP-ATG8A that co-express mCh-SidK or mCh-SidK^F62A^, treated with the tonoplast marker MDY64 displayed in magenta. Images show GFP channel, a merge of magenta and green channels, red emissions, and bright field, with and without DEX induction. Scale bar, 10 µm. **(E)** Model of the plant response to vacuolar damage upon cell wall damage through ATG8ylation of the tonoplast. Created with Biorender.com.

To identify other proteins that are involved in tonoplast ATG8ylation, we performed proximity labeling proteomics of TurboID tagged ATG8E line upon Torin, ES20, and Monensin treatments (Fig. 4B and table S1). We used TurboID alone and ATG8-AIM Docking Site mutant (ATG8^ADS^) as controls. ATG8 proxitome upon Torin treatment revealed several proteins that are known to interact with ATG8 in an AIM-dependent manner (Fig. 4B). In ES20 and Monensin treated samples, we identified several proteins that could contribute to tonoplast repair. These proteins could be grouped as vesicle trafficking, lipid storage and transport, plasma membrane, and quality control categories. For example, we identified TBC/RabGAP1 as an AIM-dependent interactor (Fig. 4B). Phylogenetic analysis revealed that TBC/RabGAP is a singleton in Arabidopsis (fig. S12). Since TBC/RabGAP protein was recently shown to coordinate lysosomal repair upon damage (*35*), we generated stable lines that co-express TBC/RabGAP1 and ATG8. Excitingly, unlike NBR1 or CFS1, TBC/RabGAP1 is recruited to the tonoplast upon cell wall damage (Fig. 4C). Further studies are necessary to functionally characterize the role of TBC/RabGAP1 and other interactors in vacuolar quality control.

Since both ES20 and monensin changed vacuolar pH and induced tonoplast ATG8ylation (Fig. 4A and fig. S10), we hypothesized that changes in vacuolar pH could be the driver of tonoplast ATG8ylation. Vacuolar pH is regulated by the V-ATPase, a molecular machine that pumps protons inside the vacuole via ATP hydrolysis (*36*). V-ATPase function is essential in plants and the available genetic mutants have pleiotropic phenotypes (*37*). To functionally probe the role of V-ATPase in tonoplast ATG8ylation, we decided to develop the Legionella effector protein SidK as a tool to study V-ATPase function in a spatiotemporally controlled manner. SidK binds to VHA-A subunit and triggers the constitutive assembly of the V_1_ and V_0_ subcomplexes, leading to the inhibition of V-ATPase activity (*38*). First, we tested whether SidK interacts with the plant V-ATPase. We expressed FLAG-tagged SidK and a SidK^F62A^ mutant that prevents binding to the V-ATPase (*38*) in *E. coli* and performed immunoprecipitation with *Arabidopsis thaliana* lysates. Western blot analysis of the pull down revealed that SidK but not SidK^F62A^ interacts with the VHA-A subunit of the V-ATPase (fig. S13A). Further mass spectrometry analysis of the immunoprecipitants also confirmed the presence of other V-ATPase subunits, demonstrating SidK as a potential tool to study V-ATPase function in plants (fig. S13B and table S2).

We then expressed SidK and SidK^F62A^ in Arabidopsis in a dexamethasone (DEX) inducible manner (fig. S13C). Lysosensor staining showed that SidK expression increased vacuolar pH, while SidK^F62A^ expression did not have a measurable effect (fig. S13D). Crucially, SidK expression mimicked ES20 and Monensin treatments and led to the colocalization of GFP-ATG8A with the tonoplast marker MDY-64 (Fig. 4D). This effect was dependent on binding to V-ATPase, since the expression of SidK^F62A^ did not change the localization of GFP-ATG8A. Altogether, these results demonstrate that SidK is a valuable tool to dissect V-ATPase function in plants, and V-ATPase plays a key role in tonoplast ATG8ylation.

## Discussion

Here, we define a conserved vacuolar quality control mechanism (VQC) that protects plant cells against the deleterious intracellular consequences of cell wall damage (Fig. 4E). In contrast to cell wall stiffening that induces autophagy, inhibition of cellulose biosynthesis or enzymatic degradation of the cell wall that mimics fungal infection triggers turgor pressure dependent ATG8ylation of the tonoplast (Figure 1). This is a clear example of Conjugation of ATG8 to Single Membranes (CASM) that has been demonstrated for various cellular compartments in mammalian cells as a stress response mediating membrane remodeling (*19, 39*). Interestingly, cell wall damage does not trigger relocation of key ESCRT proteins VPS23 or ALIX to the tonoplast (Fig. S9). Considering the well-established role of ESCRT in membrane repair (*40*), further systematic studies are necessary to test if other ESCRT proteins play a role in cell wall damage induced VQC or ESCRT dependent branches of VQC are activated upon other stress responses.

Genetic analyses of tonoplast ATG8ylation revealed that, although ATG11, an essential protein for selective autophagy, is not required; ATG8 conjugation machinery, particularly the WD40 domain of ATG16 is essential (Figure 2). Ultrastructural analysis of the *ΔCASM* line, which is only defective in tonoplast ATG8ylation, but not in autophagy, has shown that cell wall damage triggers vacuolar fragmentation and cell death (Figure 3). A previous study using yeast ATG8 has shown that ATG8 lipidation and membrane attachment could cause tubulation and budding of liposomes (*34*). ATG8ylation of tonoplast could isolate damaged compartments of the vacuole from the rest to maintain vacuolar integrity. Consistently, we see fragmented vacuoles in *ΔCASM* line (Figure 3). Further characterization of the *ΔCASM* line, together with various proteins that we identified in our proximity labeling experiments will be crucial to uncover other molecular players involved in VQC (Fig. 4B-C).

The main trigger for tonoplast ATG8ylation is likely an increase in vacuolar pH. All three inducers of tonoplast ATG8ylation – cell wall damage, ionophore treatment, and SidK expression – increase vacuolar pH (Figure 4). Similar to mammalian cells, the V-ATPase-ATG16 axis executes tonoplast ATG8ylation upon pH changes (*19, 41, 42*). However, how the plant cell senses the changes in vacuolar pH remains unknown. One of the proteins that we detected in our proximity labeling dataset is an LRR domain containing protein (Fig. 4C). LRR domain containing proteins are well known for their roles in modified-self recognition. Further studies will reveal if a similar modified-self response is activated via this LRR protein to initiate tonoplast ATG8ylation.

Moving forward, a key gap that needs to be filled is the molecular connection between cell wall integrity and VQC. The mechanical stability of a plant cell is safeguarded by intricate mechanostasis (mechanical homeostasis) pathways that balance two opposing mechanical forces: the outward turgor pressure of the cell with the inward forces of the elastic cell wall (*13*). Defects in one of the two balancing factors will destabilize the balance and threaten plant fitness. So far, the effect of cell wall damage has been studied in relation to the repair of the cell wall. How the cell adjusts the turgor pressure upon cell wall damage remained elusive. Our findings suggest a vacuolar mechanostasis pathway that promptly adjusts the turgor pressure upon cell wall damage to prevent cellular rupture (Fig. 3). Our findings suggest VQC is independent of FERONIA. A systematic analysis of the cell wall integrity sensors, particularly the cellulose integrity sensor THESEUS will be crucial to uncover the links between cell wall integrity and VQC (*43*). Further elucidation of VQC pathways will reveal the connections between VQC, the cell wall, and other cellular compartments that play a role in mechanostasis.

## Materials and Methods

### Plant material and cloning procedure

All *Arabidopsis thaliana* lines used in this study originate from the Columbia (Col0) ecotype background and are listed in Table S3. All transgenic lines were generated by floral dipping (*44*) and plasmid constructs were cloned with the GreenGate cloning method (*45*). The coding sequence of genes of interest were ordered from Twist Biosciences. A list of the plasmids produced is provided in Table S4. pATG8E::mCherry-TurboID, pATG8E::mCherry-TurboID-ATG8E and pATG8E::mCherry-TurboID-ATG8E^ADS^ lines were subjected to RT-PCR and homozygous plants with similar expression levels of TurboID were selected. For DEX-induced expression of SidK in Arabidopsis roots, full length sequence of SidK or SidK^F62A^ were synthesized (Twist) and subsequently cloned by GreenGate cloning with a C-terminal mCherry tag under control of 3xOPp into Module N. GR-LhG4 expression is driven by UBI10 promotor and cloned in Module M. Module M and N were combined in the destination vector pGGZ003 and transformed in Arabidopsis following Agrobacterium-mediated standard protocols. APEX2-ATG8A was cloned by GreenGate cloning into pGGSun with a UBI10 promoter. Homozygous plants with similar expression levels of APEX2 were selected. The knock-out mutant *atg16* and the truncation mutant ΔCASM were created by CRISPR/Cas9-mediated mutation (*46*). The plasmid pCBCDT1T2 was used as scaffold template and pHEE401E as destination vector. The guides, designed via CRISPR-P 2.0 (http://crispr.hzau.edu.cn/cgi-bin/CRISPR2/CRISPR), are GATCGGGAAACCATTGGCAT and GGTACAGGAGGAGAAAGCTA for *atg16* and GATCGGGAAACCATTGGCAT and GATCGGTTCCATCTTCATTG for *ΔCASM*. Cas9 was crossed out. ATG16 was cloned by GreenGate cloning into pGGZ003 with a UBI10 promoter. ATG16 complementation constructs were cloned by PCR from the previous vector. Each PCR (ATG16^1-195^, ATG16^1-253^, ATG16^1-295^, ATG16^1-379^) became a module which was cloned by GreenGate cloning into pGGSun with a UBI10 promoter. ATG16 and the four other delta C deletions were transformed into the *atg16* mutant background and homozygous plants were selected.

Male *Marchantia polymorpha* Takaragaike-1 (Tak-1) plants were maintained asexually and cultured through gemma on half-strength Gamborg’s B5 medium supplemented with 0.5 g/l MES, 1% sucrose and 1% agar under 50 µM/m^2^/s continuous white light at 21 °C. Plant lines used are listed in Table S2.

### Plant growth and plant treatments

For standard plant growth, seeds were sown on water-saturated soil and kept in 16h light/8h dark photoperiod with 165 µmol m-2 s-1 light intensity. For in vitro seedling growth, Arabidopsis seeds were surface sterilized in 70% ethanol 0.05% SDS for 15 minutes, rinsed in ethanol absolute and dried on sterile paper. Seeds were plated in ½ MS salts (Duchefa)/1% agar/1% sucrose plates and stratified for 48 hours in the dark at 4°C. Plates were then grown under LEDs with 50 µM/m^2^/s and a 16 h light/8 h dark photoperiod for the indicated amount of time.

For drug-treatments, all drugs used were dissolved in DMSO (unless further described) and added to the desired concentration: 9 μM Torin1 (CAS 1222998-36-8; Santa Cruz), 50 µM EGCG (E4143, Sigma-Aldrich), ES20 and ES20-1 (*16*), 3 nM Isoxaben in MS agar plates (82558-50-7, Sigma-Aldrich), 1% Driselase in PBS (85186-71-6, Sigma-Aldrich), 50 mM Sorbitol, 200 µM Monensin (22373-78-0, Sigma-Aldrich) and dexamethasone in MS agar plates (50-02-2, Sigma-Aldrich). An equal amount of pure DMSO, or respective solvent, was added to control samples.

### Carbon starvation assay

30-40 *A. thaliana* seeds were surface sterilized with ethanol, vernalized for 2 days at 4 °C in dark and grown in ½ MS media (Murashige and Skoog salt + Gamborg B5 vitamin mixture [Duchefa] supplemented with 0.5 g/liter MES and 1% sucrose, pH 5.7) for 9 days at 21°C under LEDs with 85 µM/m^2^/s with a 14 h light/10 h dark photoperiod. 9-days old seedlings were rinsed twice with new carbon-depleted ½ MS media (Murashige and Skoog salt + Gamborg B5 vitamin mixture [Duchefa] supplemented with 0.5 g/liter MES, pH 5.7) and incubated for 4 days with the carbon-depleted media in the dark. Control seedlings were rinsed twice with ½ MS media and incubated for 4 days under LEDs with 85 µM/m^2^/s with a 14 h light/10 h dark photoperiod. Pictures were taken on the 4^th^ day of treatment with a Canon EOS 80D.

### Preparation of *M. polymorpha* samples for confocal microscopy

The *M. polymorpha* asexual gemmae were incubated in liquid 1/2 Gamborg B5 media for 2 d before imaging. 2-d-old *M. polymorpha* thalli were placed on a microscope slide with deionized water and covered with a coverslip. The meristem region was used for image acquisition.

### Confocal microscopy

All images were acquired by an inverted point laser scanning confocal microscope (LSM800, Carl Zeiss) equipped with high-sensitive GaAsP (Gallium Arsenide, like LSM780 and 880) detectors, a transmitted light detector, a 20x/0.8 pan-apochromat DIC, a 63x/1.2 plan-apochromat (water immersion), and ZEN software (blue edition, Carl Zeiss). The 63x objective was used for all images except Fig. 3F, for which the 20x objective was used instead. GFP and YFP fluorescence were excited at 488 nm and detected between 488 and 545 nm. MDY64 fluorescence was excited at 405 nm and detected between 465 and 550 nm. RFP and mCherry and PI fluorescence were excited at 561 nm and detected between 570 and 617 nm. Yellow/Blue DND-160 was excited at 405 nm and detected between 400 and 500 nm for blue detection and 500 and 600 nm for yellow detection. For Z-stack imaging, interval between the layers was set as 1 μm. For each experiment, all replicate images were acquired using identical confocal microscopic parameters. Confocal images were processed with Fĳi (version 1.54, Fĳi).

Different markers were applied for 10 minutes before visualizing the samples when indicated in the figures: 50 μg/mL Propidium Iodide in PBS (25535-16-4, Sigma-Aldrich), 1 μM MDY-64 (Y7536, Invitrogen) and 20 mM Lysosensor Yellow/Blue DND-160 (L22460, Invitrogen). Mechanoprobes were also applied before measurements: 10 μM CarboTag-BDP for 30 min, 10 μM Sulfo-BDP for 30 min and 10 μM PEG-BDP for 90 min.

### Image processing and quantification

All analyses were done with 5 biological replicates per sample. Pearson’s and Spearman’s colocalization analyses were performed by Fĳi (version 1.54, Fĳi). Puncta quantification was performed by Fĳi (version 1.54, Fĳi). Z-stack images (at least five layers) were background-subtracted with 25 pixels of rolling ball radius. Each Z-stack image was subsequently thresholded using the MaxEntropy method and was converted to an 8-bit grayscale image. Threshold values were adjusted according to the puncta signals in original confocal images. The number of puncta in thresholded images was counted by the Analyze Particles function in Fĳi. For all puncta quantification, puncta with the size between 0.10 and −4.00 μm2 were counted.

### APEX2-labeling and electron microscopy sample preparation

After the respective plant treatments, the roots of seedlings were dissected (on ice) in 2.5% glutaraldehyde in 0.1 M cacodylate buffer (pH 7.4) and incubated in this fixative solution for 1h under vacuum. After washing the samples thoroughly with 0.1 M cacodylate buffer (pH 7.4; 4-5 times), the specimens were incubated in fresh prepared DAB solution with H_2_0_2_ (DAB 0.5 mg/ml, H_2_0_2_ 10 mM) in 0.1 M cacodylate buffer (*17*) for 50 min (on ice covered with tinfoil). Then, gently removing the DAB solution and washing the specimens with 0.1 M cacodylate buffer 3 times in 1 min. After that, the specimens were incubated in fresh prepared 1% (w/v) OsO_4_ in 0.1 M cacodylate buffer for 40 min at room temperature. Excess OsO_4_ was washed with deionized water (4-5 times, 15 min for each step), then gently removing the deionized water and submerging the specimens in 2% (w/v) uranyl acetate (UA) solution for 50min covered with tinfoil. Then the excess UA was washed with deionized water (4-5 times, 15 min for each step), and the specimens were dehydrated with a graduated acetone series in deionized water (from 10% to 100% acetone; 30min for each step). After embedding the samples in Embed-812 resin (Cat. No. 14120, Electron Microscopy Sciences) and polymerized at 60 °C oven for 24 h. The ultrathin sections (70 nm thick) were prepared from the sample blocks. The sections were examined with a transmission electron microscope (Morgagni 268) operated at 80 kV.

### Observation of tonoplast by transmission electron microscopy (TEM) and Electron tomography (ET)

After the respective plant treatments, the roots of seedlings were dissected (on ice) in 2.5% glutaraldehyde in 0.1 M cacodylate buffer (pH 7.4) and incubated in this fixative solution overnight at 4 °C. After washing the samples thoroughly with 0.1 M cacodylate buffer (pH 7.4; 4-5 times), the specimens were postfixed by incubation in fresh prepared 1% (w/v) OsO_4_ in 0.1 M cacodylate buffer for 1h at room temperature. Excess OsO_4_ was washed with deionized water (4-5 times, 15 min for each step), and the samples were dehydrated with a graduated acetone series in deionized water (from 10% to 100% acetone; 30min for each step). After embedding the samples in Embed-812 resin and polymerized at 60 °C oven for 24 h. Thin sections (90 nm thick) were prepared from the sample blocks. The sections were poststained and examined with a transmission electron microscope (Morgagni 268) operated at 80 kV.

A series of semi-thick sections (250 nm) were collected on copper slot grid (Cat. No. GS2010-Cu, Electron Microscopy Sciences). After post-staining and gold particle coating, tilt series were collected with a 200-kV Tecnai G2 20 electron microscope (+ 50° to −50° at an interval of 1° interval around two orthogonal axes). Tomogram calculation and 3D model rendering were performed with the IMOD software package as described previously (*47, 48*).

### Fluorescence lifetime imaging (FLIM) microscopy

FLIM imaging experiments were performed on a Picoquant Fluorescent Lifetime Imaging Microscope equipped with Olympus iX71 inverted microscope frame, PL 20 x PlanAchromat Objektiv, NA = 0.4, NA = 1.2 (water immersion), Hybrid Photomultiplier Detection Assembly with < 50 ps time resolution and SymPhoTime 64 (Picoquant). Samples were excited with a 488-nm pulsed laser source (pulse duration <1 ps). Acquisition time was fixed at 120 s for each 256 × 256 pixel image. FLIM images were processed using SymPhoTime 64 software to fit the fluorescence decay curves in each pixel with a two-component exponential decay. Images are reported in a false-color scale that represent the mean fluorescence lifetime for each pixel, expressed in nanoseconds. 3 technical replicates were measured by session and their average is represented in the final graph. 3 biological replicates were measured in different days and its average mean value is provided as the final value.

### Protein extraction and western blotting

20-40 *A. thaliana* seeds were surface sterilized with ethanol, vernalized for 2 days at 4 °C in dark and grown in ½ MS media (Murashige and Skoog salt + Gamborg B5 vitamin mixture [Duchefa] supplemented with 0.5 g/liter MES and 1% sucrose, pH 5.7) for 7 days at 21°C under LEDs with 85 µM/m^2^/s with a 14 h light/10 h dark photoperiod. For drug treatments, Monensin (CAS 22373-78-0; Sigma-Aldrich) was dissolved in pure ethanol, Torin1 (CAS 1222998-36-8; Santa Cruz) and ES20 (*49*) in DMSO, then added to the desired concentration: 200 mM Monensin, 3 μM Torin1, 500 μM ES20. Equal amounts of pure ethanol or DMSO were added to mock samples. Seedlings were harvested in safe lock Eppendorf tubes containing 2 mm Ø glass beads, flash frozen in liquid nitrogen and ground using a Silamat S7 (Ivoclar vivadent). Total proteins were extracted in 2X Laemmli buffer by shaking again in the Silamat S7 for 20 s. Samples were boiled for 10 min at 70 °C and 1000 rpm shaking, then centrifuged for 5 min at maximum speed. Total proteins were quantified with the amido black method. 10 μl of supernatant was mixed with 190 μl of deionized water and added to 1 ml of Amido Black Buffer (10% acetic acid, 90% methanol, 0.05% [w/v] Amido Black (Naphtol Blue Black, Sigma N3393)), mixed and centrifuged for 10 min at maximum speed.

Pellets were then washed with 1 ml of Wash Buffer (10% acetic acid, 90% ethanol), mixed and centrifuged for 10 min at maximum speed and resuspended in 0.2N NaOH. OD_630 nm_ was measured, with NaOH solution as blank, and protein concentration was calculated using the OD = *a*[C] + *b* determined curve. 2.5–40 μg of total protein extracts were separated on SDS-PAGE gels and blotted onto PVDF Immobilon-P membrane (Millipore). GFP was detected using the anti-GFP antibody (Mouse monoclonal, 11814460001; Roche) diluted 1:5000 (v/v). NBR1 was detected using the anti-NBR1 antibody (Rabbit polyclonal, AS14 2805; Agrisera). The subunit A of the V-ATPase was detected using the anti-V-ATPase-A antibody (Rabbit polyclonal). Mouse monoclonal antibodies were detected with goat anti-mouse IgG HRP-linked antibody (61-6520; Invitrogen) diluted 1:5000 (v/v). Rabbit polyclonal antibody was detected with a goat anti-rabbit IgG HRP-linked antibody (65-6120; Invitrogen) diluted 1:5000. Hybridized membranes were reacted with SuperSignal™ West Pico PLUS Chemiluminescent Substrate (Thermo Fisher Scientific) and imaged using an iBright CL1500 Imaging System (Invitrogen).

### Expression and purification of SidK in *E. coli*

For protein production in E. coli, amino acid residues Tyr9 to Asp278 from SidK WT and SidK^F62A^ were amplified and subsequently cloned into pOPIN-GG vector pPGN-C (Bentham et al. 2021) with a N-terminal 6xHIS-3C and a C-terminal 3xFLAG via Golden Gate cloning (*50*).

Proteins were produced following established protocols for effector purification (*51*). Briefly, plasmids were transformed into *E. coli* RosettaTM (DE3) and a single colony was grown overnight in LB media supplemented with appropriate antibiotics. Cell cultures were then grown in terrific broth media at 37°C for 5–7 hr and then at 16°C overnight. Cells were harvested by centrifugation and re-suspended in 50 mM Tris-HCl (pH 8), 500 mM NaCl, 5% (vol/vol) glycerol, and 20 mM imidazole supplemented with EDTA-free protease inhibitor tablets (Roche). Cells were sonicated and following centrifugation at 40,000xg for 30 min, the clarified lysate was applied to a HisTrapTM Ni2+-NTA column connected to an ÄKTA Pure chromatography system (Cytiva Life Sciences). Proteins were step-eluted with elution buffer (50 mM Tris-HCl (pH 8), 500 mM NaCl, 5% (vol/vol) glycerol, and 500 mM imidazole) and directly injected onto a Superdex 200 16/60 gel filtration column pre-equilibrated with 20 mM HEPES (pH 8), 150 mM NaCl and 5% (vol/vol) glycerol supplemented with 1mM TCEP. Elution fractions were collected and evaluated by SDS-PAGE. Relevant fractions were pooled together and concentrated as appropriate.

### Pull down of plant V-ATPase from plant extracts

To pull down native V-ATPase from plant extracts, Arabidopsis leaf tissue was collected and ground to fine powder in liquid nitrogen using a pestle and mortar. Leaf powder was mixed with two times volume/weight ice-cold extraction buffer (10% glycerol, 25 mM Tris pH 7.5, 1 mM EDTA, 150 mM NaCl, 2% w/v PVPP, 10 mM DTT, 0.2% IGEPAL® [Merck]) supplemented with EDTA-free protease inhibitor tablets (Roche), centrifuged at 5000x g at 4°C for 20–30 min, and the supernatant was passed through a 0.45 μm Minisart syringe filter.

For immunoprecipitation, 6xHis:GFP:3xFLAG, 6xHis:SidK:3xFLAG or 6xHis:SidKF62A:3xFLAG protein were added to 1 ml of filtered extract to a final concentration of ∼2.5 μM and incubated in a rotatory mixer at 4°C with 25 μl of Anti-FLAG® M2 Magnetic Beads (Merck). 30 μl of the mixture was taken as input for Western blot analysis before adding magnetic beads. After 2.5 hr the beads were pelleted using a magnetic rack and the supernatant removed. The pellet was washed by resuspension in 1 ml of IP buffer (10% glycerol, 25 mM Tris pH 7.5, 1 mM EDTA, 150 mM NaCl, 0.2% IGEPAL® [Merck]) and pelleted again in the magnetic rack. Washing steps were repeated three times with IP buffer and three times with 50 mM Tris pH 7.5, 150 mM NaCl.

Finally, the beads were pelleted by centrifugation and incubated for 10 min at 70°C with SDS loading buffer. The beads were then pelleted again, and the supernatant loaded on SDS-PAGE gels prior to western blotting. Membranes were probed with anti-FLAG M2 antibody (Merck) to detect SidK and anti-A (AS09 467, Agrisera) to detect A subunit of the V-ATPase.

### *In vivo* co-immunoprecipitation

For co-immunoprecipitation, 14 mg of seeds per sample were grown in ½ MS media for 7 days. Proteins were extracted by adding 3 equivalent volumes of Extraction Buffer (10% glycerol, 10 mM DTT, 10 mM Tris pH 7.4, 50 mM KCl, 5 mM MgCl2, 5mM ATP, 0.5% dodecyl beta-D-maltoside, 20 μg/mL Pepstatin, 1 tablet/50 mL cOmplete EDTA-free Protease Inhibitor Cocktail [Roche]). Lysates were cleared by centrifugation at 4000 rpm at 5°C for 15 minutes three times. After the first centrifugation, the supernatant was filtered with Miracloth (Sigma-Aldrich). The supernatant was incubated with 30 μl GFP-Trap ® Agarose beads (Chromotek) for 1,5 h. Beads were washed three times with Wash Buffer (10% glycerol, 10 mM DTT, Tris pH 7.4, 50 mM KCl, 2mM MgCl2, 1 mM ATP, 0.1% dodecyl beta-D-maltoside, 20 μg/mL Pepstatin, 1 tablet/50 mL cOmplete EDTA-free Protease Inhibitor Cocktail [Roche]) before and after incubation with lysate. Beads were eluted in 100 μl 2x Laemmli buffer, boiled for 5 min at 95 °C and subjected to Western blot with indicated antibodies.

### ES20-1 synthetic procedure

o-Methyl benzoyl hydrazine (3.00 g, 20 mmol, 1.00 equiv.) was dissolved in 80 mL abs. ethanol under stirring and benzoyl isothiocyanate (3.26 g, 20 mmol, 1.00 equiv.) was added. After some minutes of stirring, a precipitate formed. The mixture was heated to reflux for 15 minutes. The solution was then allowed to cool to room temperature, whereas colorless crystals started to form. To complete crystallization, the flask containing the crystals and the mother liquor was cooled in an ice bath. The product was filtered, washed with cold ethanol and dried in vacuo. The product (4.90 g, 15.6 mmol, 78 % yield) was obtained as an off-white crystalline solid.

^1^H NMR (700 MHz, CDCl3): δ= 13.45 (s, 1H), 9.48 (s, 1H), 9.05 (s, 1H,), 7.92-7.88 (m, 2H), 7,69-7.62 (m, 1H), 7.59 (d, 1H, *J* = 7.55Hz), 7.56-7.51 (m, 2H), 7.45-7.39 (m, 1H), 7.31-7.27 (m, 2H), 2.56 (s, 3H) ppm.

^13^C NMR (176 MHz, CDCl3) δ= 171.35, 166.56, 164.46, 137.87, 133.83, 131.60, 131.46, 131.40, 131.06, 129.21, 127.61, 127.53, 126.03, 20.19 ppm.

HRMS (ESI): m/z calcd. for [C16H15N3O2S, M+Na]+: 336.0777; found: 336.0767.

### Phylogenetic analysis of TBC-RABGAP1

To build a phylogeny of TBC-RABGAP1, we first use BLASTP from BLAST+ suite (*52*) to search for sequences closely related to AT5G52580.1 in The Arabidopsis Information Resource (TAIR) database, Solanaceae Genomics Network (https://solgenomics.net; genomes: Niben101 and Capang) and Phytozome (https://phytozome-next.jgi.doe.gov; genomes: A.thaliana_Araport11, A.lyratav2.1, C.rubellav1.1, E.salsugineumv1.0, T.cacaov2.1, P.vulgarisv2.1, G.maxWm82.a4.v1, M.truncatulaMt4.0v1, L.japonicusLj1.0v1, S.lycopersicumITAG5.0, S.tuberosumv6.1, A.comosusv3, A.trichopodav1.0, P.virgatumv5.1, S.bicolorv3.1.1, Z.maysRefGen_V4, O.sativav7.0, H.vulgare_MorexV3, T.aestivumv2.2, B.distachyonv3.1, M.polymorphav3.1, S.moellendorffiiv1.0, C.reinhardtii_CC-4532v6.1 and P.patensv3.3). In total we collected 49 non-redundant sequences from 26 species (Supplemental dataset 1). Amino acid-based alignment was generated using MUSCLE (*53*) and was subsequently trimmed from poorly aligned positions using Gblocks (*54*) with less stringent parameter as implemented in http://phylogeny.lirmm.fr/phylo_cgi/. The resulting blocks were used to compute a Maximum Likelihood phylogenetic tree using IQ-Tree 2 (*55*). Best-scoring tree was visualized using the iToL tool 6.9 (*56*) and is publicly available at https://itol.embl.de/export/1931711883132731712669776.

### Affinity purification of biotinylated proteins and nanoLC-MS/MS Analysis

*A. thaliana* seeds were surface sterilized with ethanol, stratified for 2 days and grown in ½ MS (Duchefa)/0.5% MES/1% sucrose for 7 days under LEDs with 50 µmol m-2 s-1 and a 16 h light/8 h dark photoperiod. 7-days old seedlings were incubated with 50 µM biotin for 4 hours contextually with ES20 or Torin1 treatment, whereas Monensin-treated seedlings were incubated with 100 µM biotin for 2 hours. After the treatment, the seedlings were quickly rinsed in ice cold water, dried and frozen in liquid nitrogen. For the affinity purification of biotinylated proteins, around 1 gram of plant tissue was used for each sample and the protocol was performed as described by Mair et al. (2019) (*57*).

For MS Analysis, the nano HPLC system (UltiMate 3000 RSLC nano system) was coupled to an Orbitrap Exploris 480 mass spectrometer equipped with a Nanospray Flex ion source for the Exploris 480 (all parts Thermo Fisher Scientific). Peptides were loaded onto a trap column (PepMap Acclaim C18, 5 mm × 300 µm ID, 5 µm particles, 100 Å pore size, Thermo Fisher Scientific) at a flow rate of 25 µl/min using 0.1% TFA as mobile phase. After loading, the trap column was switched in line with the analytical column (PepMap Acclaim C18, 500 mm × 75 µm ID, 2 µm, 100 Å, Thermo Fisher Scientific). Peptides were eluted using a flow rate of 230 nl/min, starting with the mobile phases 98% A (0.1% formic acid in water) and 2% B (80% acetonitrile, 0.1% formic acid) and linearly increasing to 35% B over the next 120 min. This was followed by a steep gradient to 95%B in 1 min, stayed there for 6 min and ramped down in 2 min to the starting conditions of 98% A and 2% B for equilibration at 30°C. The Orbitrap Exploris 480 mass spectrometer was operated in data-dependent mode ‘Cycle Time’, performing a full scan (m/z range 350-1200, resolution 60,000, normalized AGC target 300%, compensation voltages CV of -45V, -60V and -75V), followed by MS/MS scans of the most abundant ions for a cycle time of 0.9 seconds per CV. MS/MS spectra were acquired using an isolation width of 1.2 m/z, normalized AGC target 200%, HCD collision energy of 30, orbitrap resolution of 30,000, maximum injection time of 100 ms and minimum intensity set to 25,000. Precursor ions selected for fragmentation (include charge state 2-6) were excluded for 45 s. The monoisotopic precursor selection (MIPS) mode was set to Peptide and the exclude isotopes feature was enabled.

### MS Data processing

For peptide identification, the RAW-files were loaded into Proteome Discoverer (version 2.5.0.400, Thermo Scientific). All MS/MS spectra were searched using MSAmanda v2.0.0.19924 (*58*). The peptide mass tolerance was set to ±10 ppm and fragment mass tolerance to ±10 ppm, the maximum number of missed cleavages was set to 2, using tryptic enzymatic specificity without proline restriction. Peptide and protein identification was performed in two steps. For an initial search the RAW-files were searched against the Arabidopsis database called TAIR10 (32,785 sequences; 14,482,855 residues), supplemented with common contaminants and sequences of tagged proteins of interest using beta-methylthiolation on cysteine as a fixed modification. The result was filtered to 1 % FDR on protein level using the Percolator algorithm (*59*) integrated in Proteome Discoverer. A sub-database of proteins identified in this search was generated for further processing. For the second search, the RAW-files were searched against the created sub-database using the same settings as above and considering the following additional variable modifications: oxidation on methionine, deamidation on asparagine and glutamine, phosphorylation on serine, threonine and tyrosine, biotinylation on lysine, ubiquitinylation residue on lysine, glutamine to pyro-glutamate conversion at peptide N-terminal glutamine and acetylation on protein N-terminus. The localization of the post-translational modification sites within the peptides was performed with the tool ptmRS, based on the tool phosphoRS (*60*). Identifications were filtered again to 1 % FDR on protein and PSM level, additionally an Amanda score cut-off of at least 150 was applied. Proteins were filtered to be identified by a minimum of 2 PSMs in at least 1 sample. Protein areas have been computed in IMP-apQuant (*61*) by summing up unique and razor peptides. Resulting protein areas were normalised using iBAQ (*62*) and sum normalisation was applied for normalisation between samples.

For quality control, proteins with PSMs >= 5 were filtered for each Torin-treated sample. Known ATG8 interactors specifically enriched in TID-ATG8E Torin sample are shown in Fig. 4B. Proteins with log2FC >=1 in TID-ATG8 and TID-ATG8^ADS^ samples, respectively to the correspondent treatment in TID control, were filtered to identify AIM-dependent ATG8 interactors upon ES20 and Monensin treatment, shown in Fig. 4B. Venn diagrams were produced with eulerr R package version 6.1.1 (https://CRAN.R-project.org/package=eulerr).

## Acknowledgements

We thank Vienna Biocenter Core Facilities (VBCF), particularly Proteomics, BioOptics, Electron Microscopy, and Plant Sciences. We acknowledge funding from Austrian Academy of Sciences, Austrian Science Fund (FWF, P32355, P34944, ESP 580), Austrian Science Fund (FWF-SFB F79), Vienna Science and Technology Fund (WWTF, LS17-047, LS21-009), European Research Council Grant (Project number: 101043370), Vienna International Postdoctoral Program (VIP2) and Marie Curie Fellowship to JJ and JC (Project number: 847548) and MS (Project number: 101107472). MB and JS are funded by the European Research Council (ERC CoG Catch, project number: 101000981). We thank Alberto Moreno Cencerrado for his help with the FLIM microscope. We thank Suayib Üstün, Karin Schumacher, Jürgen Kleine-Vehn, Niko Geldner, Daniel Hofius, Erika Isono, Liwen Jiang, Takashi Ueda, Alyona Minina and Vicente Rubio for providing Arabidopsis and Marchantia lines. We thank Eva Sophie Wallner and Liam Dolan for providing vectors for DEX inducible system. We thank Lei Huang for providing ES20.

## Author contributions

Conceptualization: Y.D., J.J. Methodology: J.J., P.G., A.D.C., J.C.D.L.C., M.S., M.C., M.B., C.D., N.S.C., Y.D. Investigation: J.J., P.G., A.D.C., J.C.D.L.C., L.A., M.S., H.D., M.C., N.G., R.K., I.P. Visualization: J.J., A.D.C., P.G., J.C.D.L.C. Funding acquisition: Y.D., J.J., A.B., B.K. Project administration: Y.D. Supervision: C.D., B.K., A.B., N.S.C., J.S., Y.D. Writing – original draft: J.J., P.G., J.C.D.L.C., Y.D. Writing – review & editing: J.J., P.G., A.D.C., J.C.D.L.C., L.A., H.D., R.K., C.D., A.B., N.S.C., J.S., Y.D.

## Data availability

All the source data used to generate the main and supplementary figures are deposited to Zenodo (10.5281/zenodo.10993280).

---

**Figure S1.**
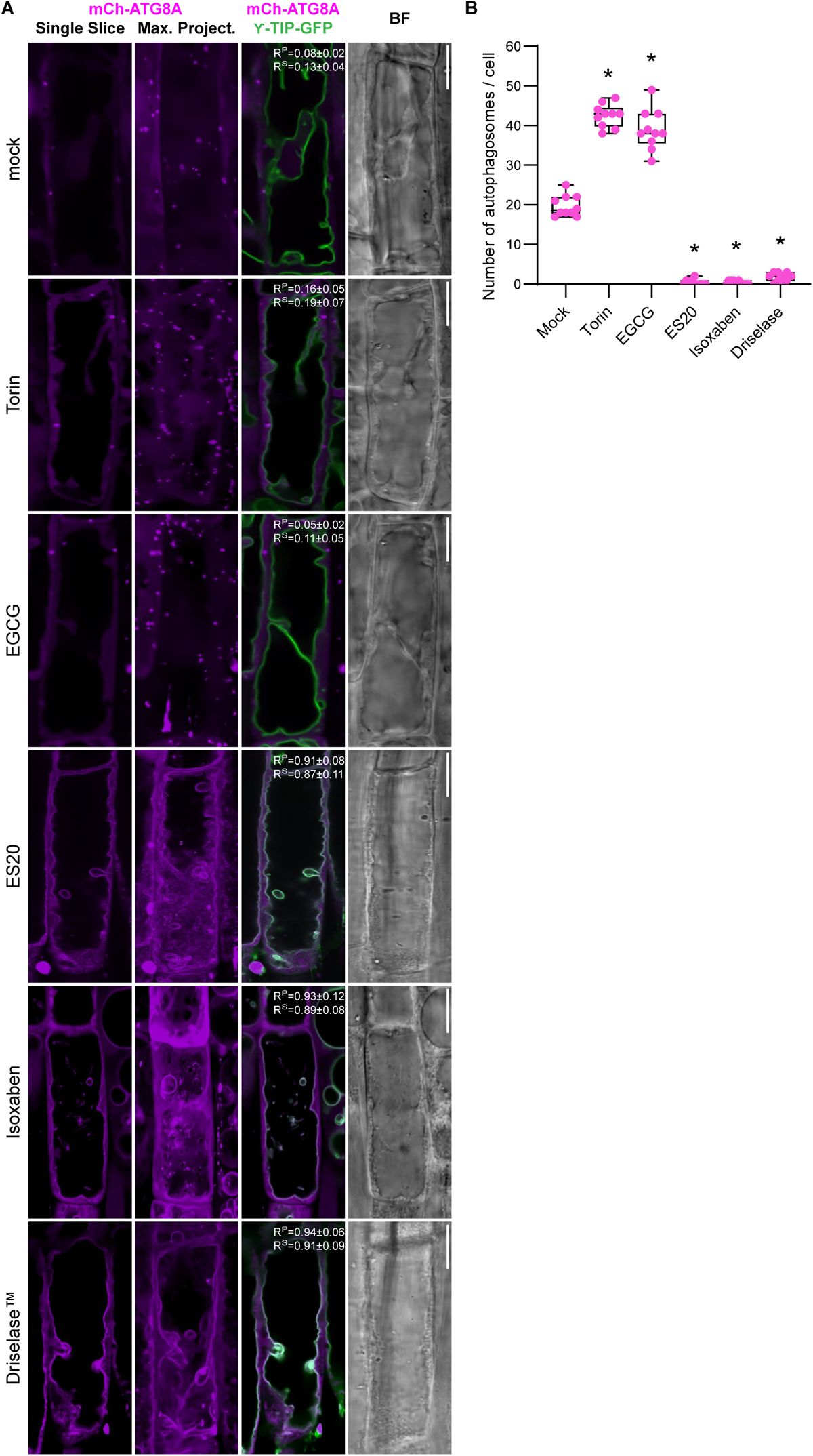
Cell wall damage triggers ATG8ylation of the tonoplast. **(A)** Confocal micrographs of the early elongation zone root cells of *Arabidopsis thaliana*, depicting the co-localization of mCherry-ATG8A (in magenta) with the tonoplast marker γ-tip-GFP. The set includes a single optical slice and a maximum intensity projection of an entire cell (20 µm depth), along with a merged image incorporating γ-tip-GFP and a corresponding bright field image. Scale bar, 10 µm. Pearson and Spearman co-localization scores are provided to quantify the association between ATG8A and the tonoplast, under treatment conditions including mock, Torin (1.5 hours, 9 µM), EGCG (30 minutes, 50 µM), ES20 (8 hours, 100 µM), Isoxaben (3 days, 3 nM), and Driselase (1 hour, 1%). **(B)** Quantitative analysis of autophagosome numbers under the treatment conditions shown in Fig. S1A. Statistical analysis was performed using Wilcoxon test to compare each treatment condition against the mock, indicating significant differences at p-values lower than 0.01.

**Figure S2.**
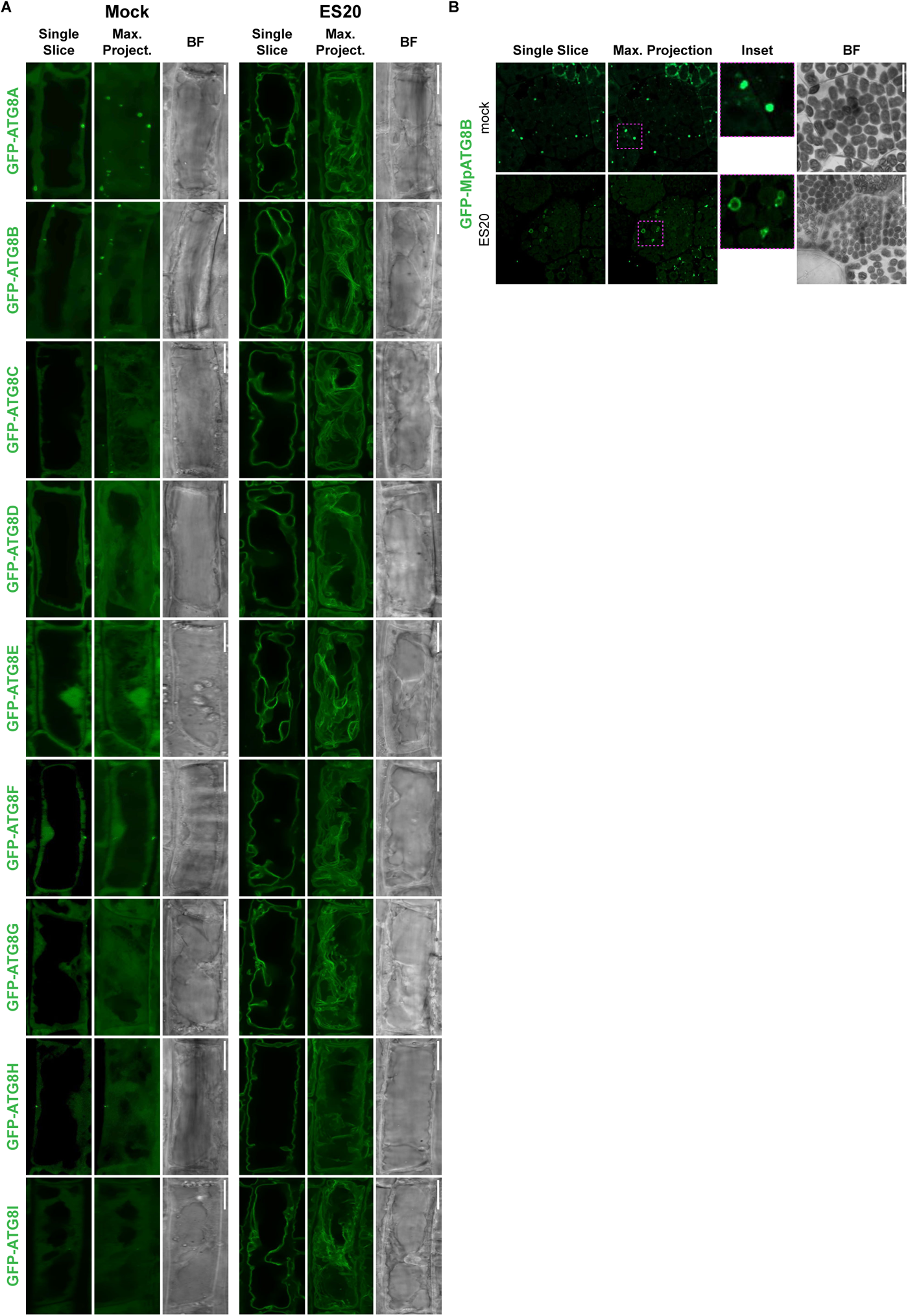
All Arabidopsis and Marchantia ATG8 isoforms are recruited to the tonoplast upon cell wall damage. **(A)** Confocal micrographs displaying the localization of all nine GFP-tagged ATG8 isoforms (GFP-ATG8A to GFP-ATG8I) in *Arabidopsis thaliana* root cells. For each isoform, images under mock conditions include a single optical slice, a maximum intensity projection of an entire cell (20 µm depth), and a corresponding bright field image. The same set of images is presented for cells treated with ES20 (8 hours, 100 µM). Scale bar, 10 µm. **(B)** Confocal micrographs of *Marchantia polymorpha*, comparing GFP-ATG8B localization under mock and ES20 (8 hours, 100 µM) treatment conditions. Scale bar, 10 µm.

**Figure S3.**
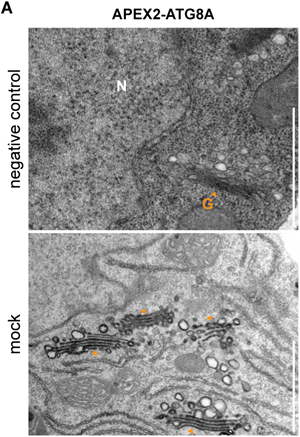
Background Labeling in APEX2-ATG8A samples. **(A)** Electron microscopy (EM) images displaying APEX2-ATG8A localization post DAB staining to highlight unspecific labeling of the Golgi stacks in APEX2-ATG8A lines. N: Nucleus, G and orange arrowheads: Golgi apparatus. Scale bar, 1 µm.

**Figure S4.**
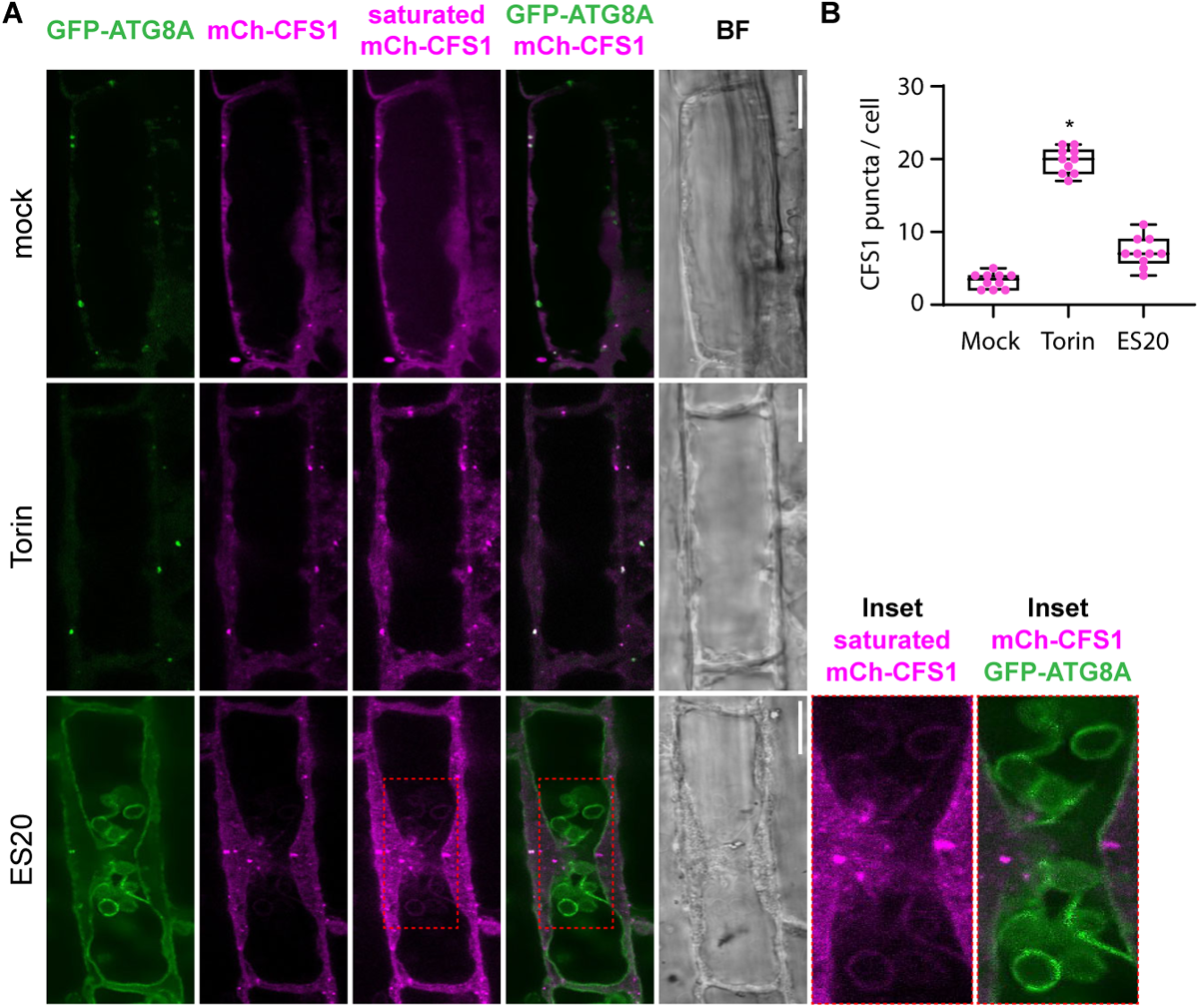
Autophagy adaptor CFS1 is not recruited to the tonoplast upon cell wall damage. **(A)** Confocal micrographs obtained from GFP-ATG8A mCh-CFS1 stably co-expressing *Arabidopsis thaliana* root cells under mock, Torin (1.5 hours, 9 µM), and ES20 (8 hours, 100 µM) treatments. The panel sequence includes single optical slices for GFP-ATG8A and mCh-CFS1, an additional panel for mCh-CFS1 with oversaturation to enhance visualization, a merge of both channels, corresponding bright field images and insets to enhance visualization. Scale bar, 10 µm. **(B)** Quantification of CFS1 puncta across the different treatment conditions shown in Fig. S4A. Statistical analysis was performed using Wilcoxon test to compare each treatment condition against the mock, indicating significant differences at p-values lower than 0.01.

**Figure S5.**
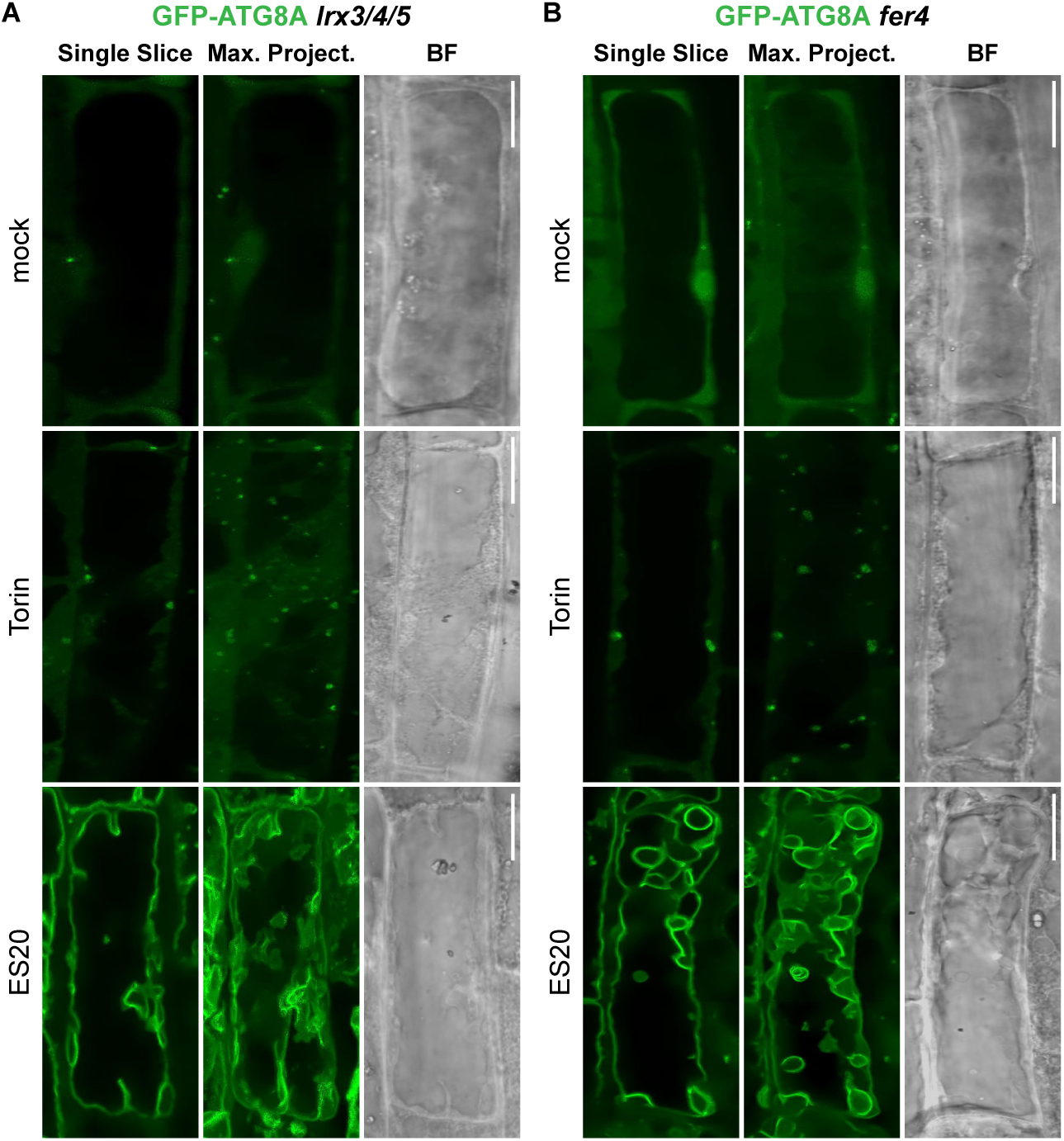
Tonoplast ATG8ylation is not regulated by FERONIA. **(A)** Confocal micrographs of GFP-ATG8A expressed in *Arabidopsis thaliana lrx3/4/5* triple mutant background **(A)** or *fer4* mutant background **(B)**, treated with mock, Torin (1.5 hours, 9 µM), and ES20 (8 hours, 100 µM). The set includes a single optical slice, a maximum intensity projection, and a bright field image for each treatment condition. Scale bar, 10 µm.

**Figure S6.**
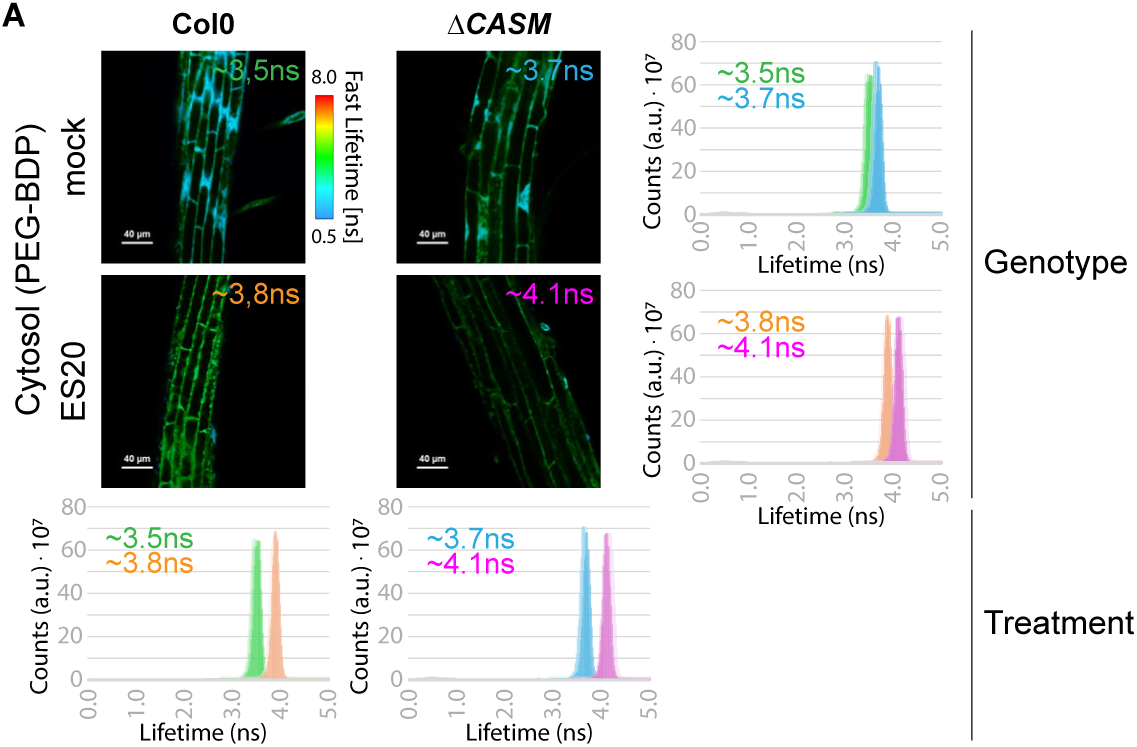
FLIM analysis of PEG-BDP mechanoprobe to assess cytoplasmic crowding. Analysis of Col0 and *ΔCASM* backgrounds treated with PEG-BDP, a fluorescent cytoplasmic mechanoprobe, for 90 minutes at 10 µM concentration, following mock or ES20 (8 hours, 100 µM) treatments. The fluorescence lifetime of the probe across three biological replicates, with average lifetimes reported in nanoseconds. Four comparative graphs detail the lifetime variance per treatment and genotype. Scale bar, 40 µm.

**Figure S7.**
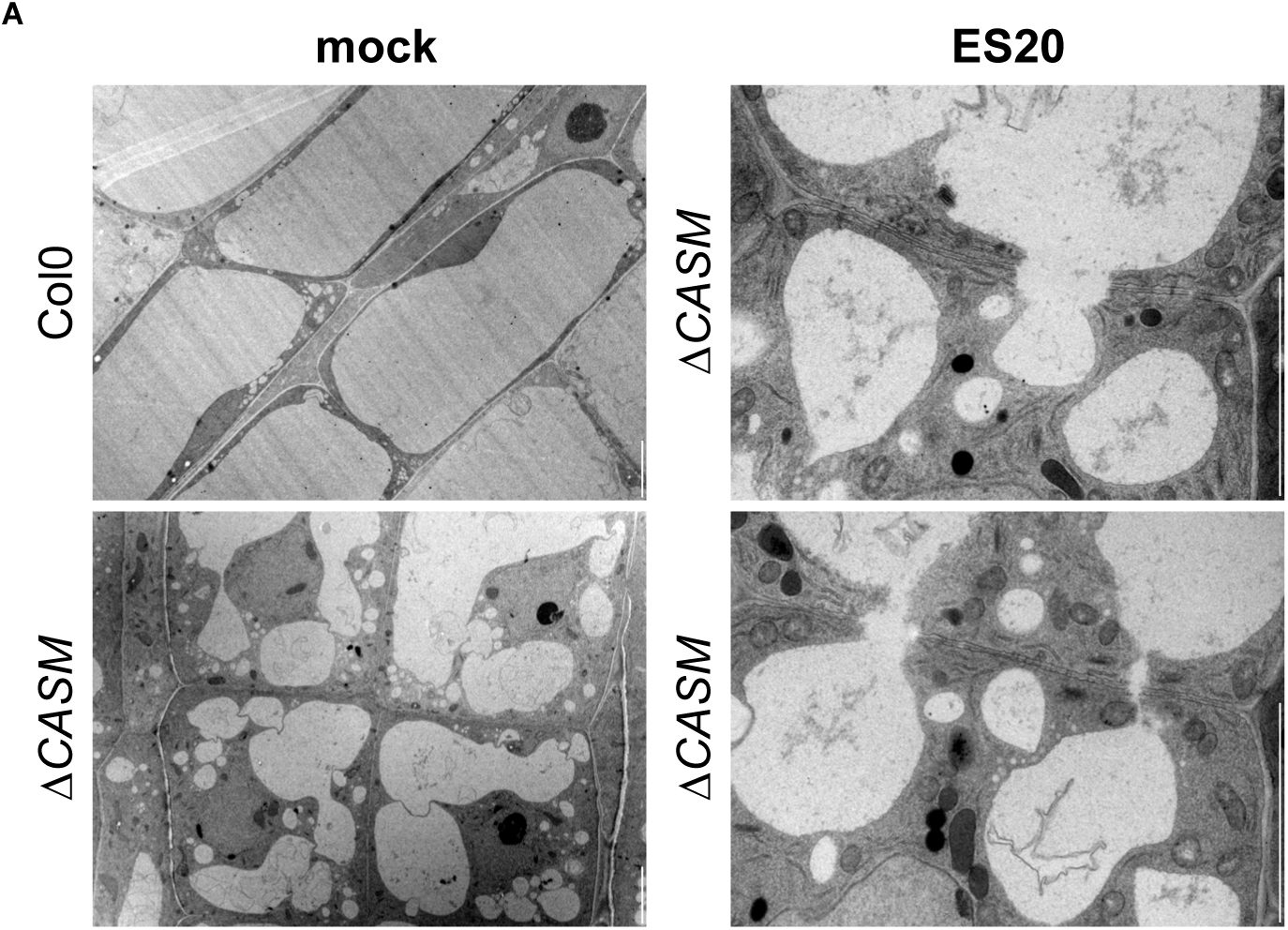
Vacuolar morphology of *ΔCASM* root cells upon cell wall damage. Transmission Electron Microscopy (EM) images visualizing the vacuole in Col0 and *ΔCASM* backgrounds under mock and ES20 (8 hours, 100 µM) treatments. The images reveal vacuolar fusion between cells in response to cell wall damage. Scale bar, 5 µm.

**Figure S8.**
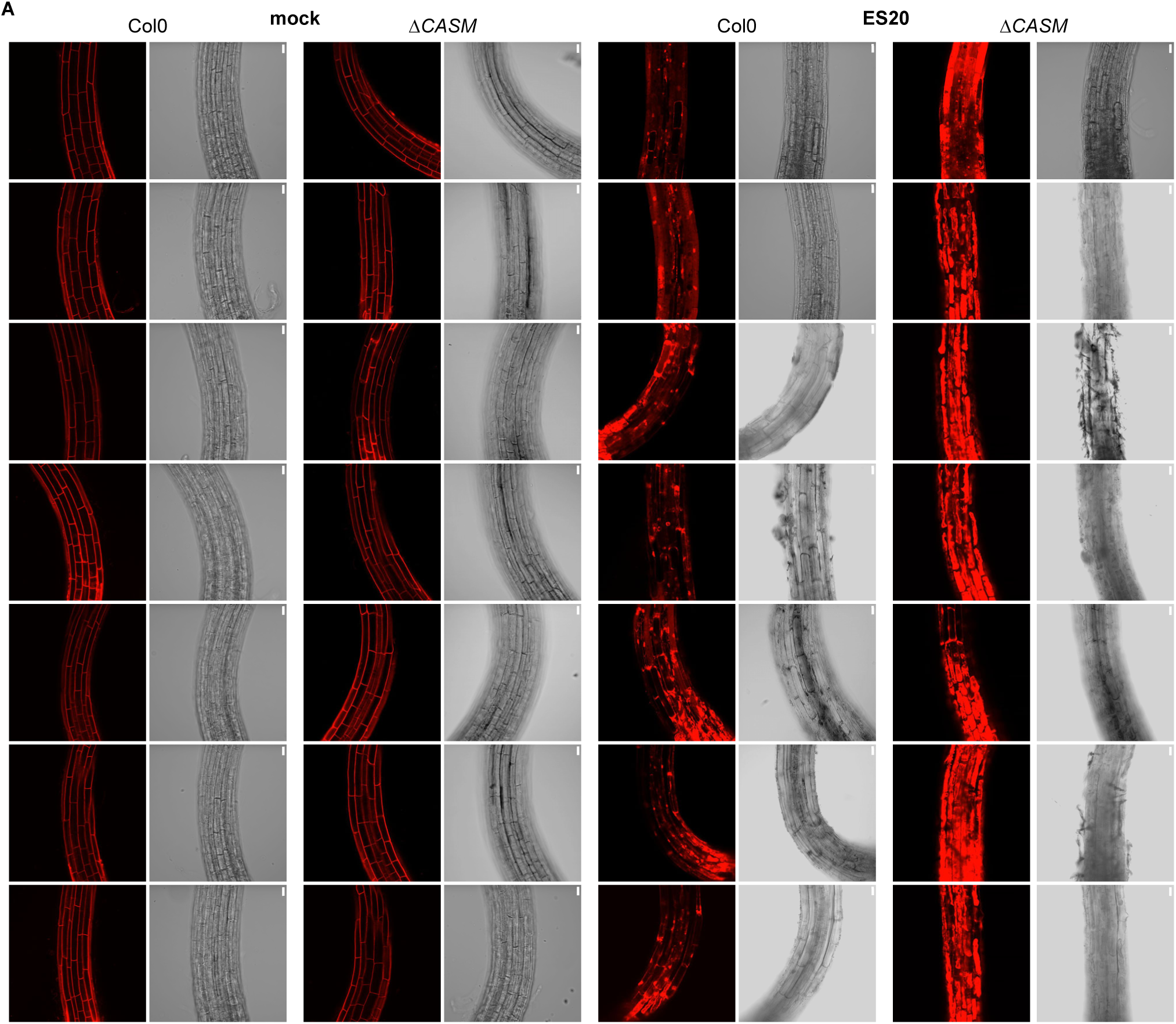
*ΔCASM* root cells are more sensitive to cell wall damage. Propidium Iodide (PI) staining of root cells from Col0 and *ΔCASM* backgrounds under mock and ES20 (8 hours, 100 µM) treatments, assessing cell viability and membrane integrity. Seven more replicates of Fig. 3F are shown for each genotype and treatment. Scale bar, 10 µm.

**Figure S9.**
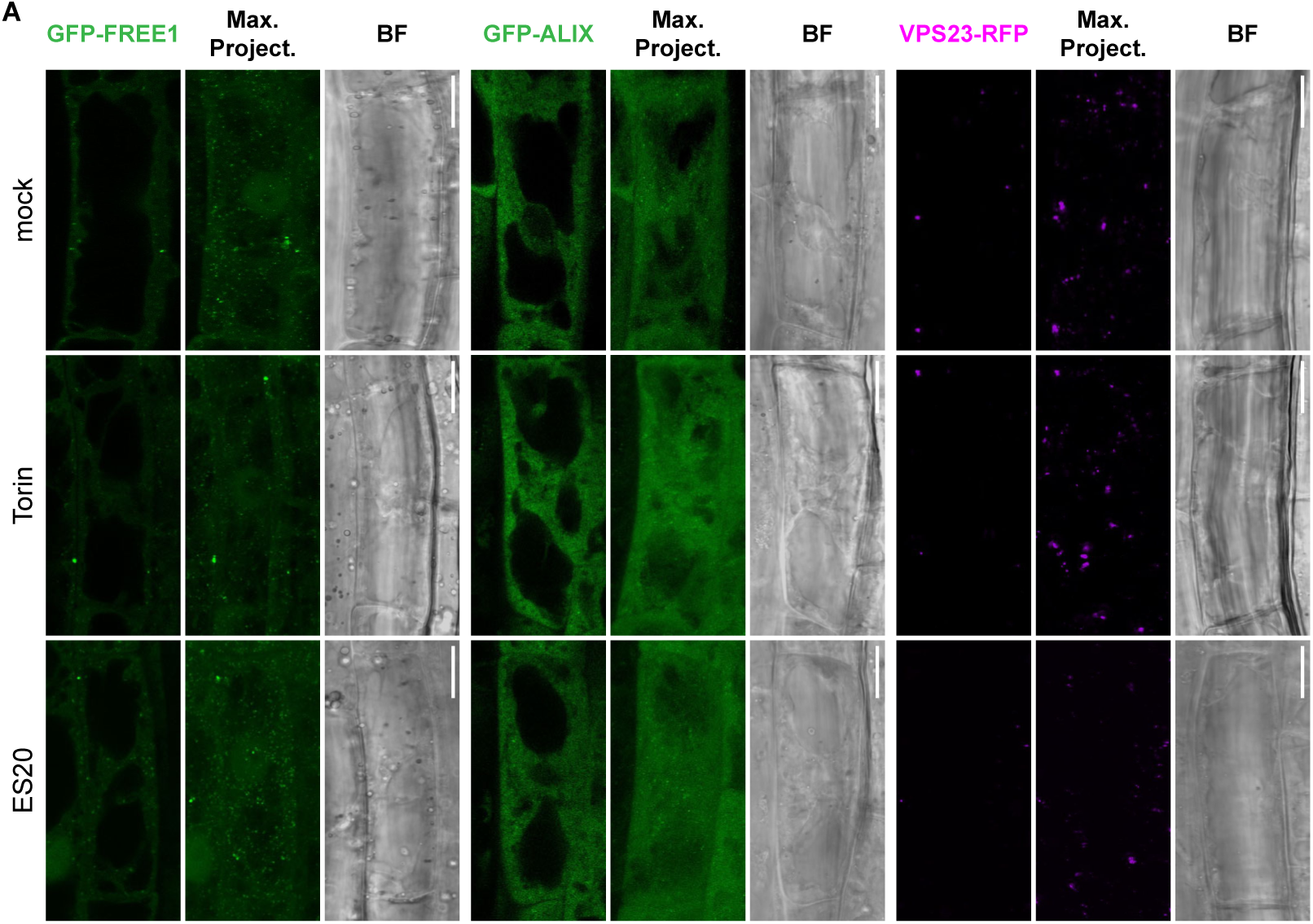
Key ESCRT proteins are not recruited to the tonoplast upon cell wall damage. Confocal micrographs depicting the localization of ESCRT (Endosomal Sorting Complex Required for Transport) machinery components GFP-FREE1, GFP-ALIX, and VPS23-RFP in *Arabidopsis thaliana* root cells, under mock, Torin (1.5 hours, 9 µM), and ES20 (8 hours, 100 µM) treatments. Each set comprises a single optical slice, a maximum intensity projection, and a corresponding bright field image, facilitating a comparative analysis of ESCRT component dynamics under canonical and non-canonical autophagy inducing conditions. Scale bar, 10 µm.

**Figure S10.**
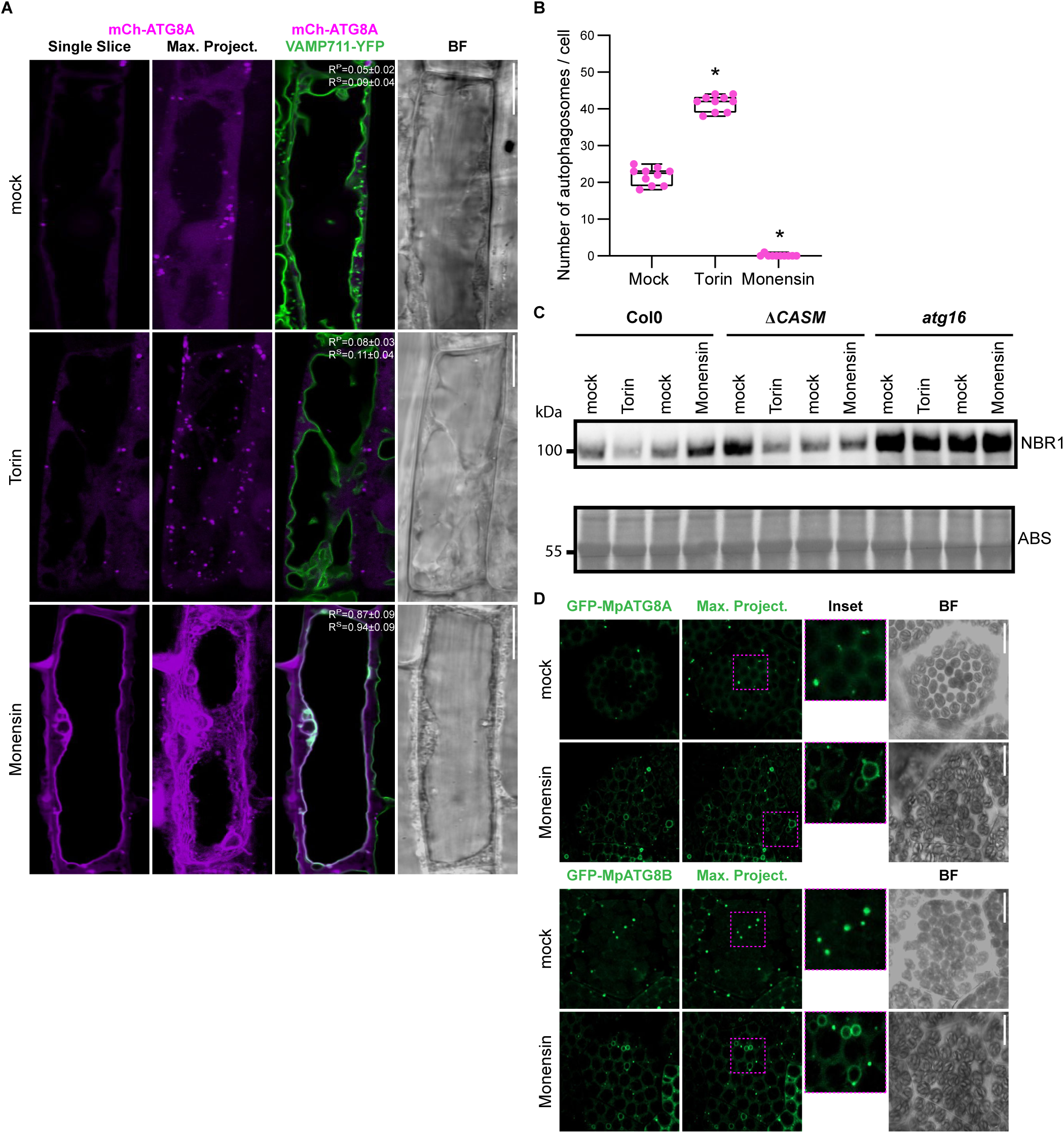
Vacuolar ionophore monensin triggers tonoplast ATG8ylation. **(A)** Confocal micrographs of root cells in the early elongation zone of *Arabidopsis thaliana*, highlighting the localization of mCherry-ATG8A (in magenta) illustrating tonoplast ATG8ylation. The panel includes a single optical slice and a maximum intensity projection of a whole cell (20 µm depth), alongside a merged image with VAMP711-YFP (tonoplast marker) and a corresponding bright field image. Scale bar, 10 µm. Pearson and Spearman co-localization values are presented, showing the association between ATG8A and the tonoplast. Treatment conditions include mock, Torin (1.5 hours, 9 µM), and Monensin (0.5 hours, 200 µM) treatments. **(B)** Quantification of autophagosomes under various treatment conditions depicted in Fig S9A. Statistical analysis was performed using Wilcoxon test to compare each treatment condition against the mock, indicating significant differences at p-values lower than 0.01. **(C)** Western blot analysis of NBR1 flux under mock, Torin (1.5 hours, 9 µM), and Monensin (0.5 hours, 200 µM) treatments. **(D)** Confocal micrographs of GFP-MpATG8A and GFP-MpATG8B expressing *Marchantia polymorpha* cells under mock, Torin (4 hours, 9 µM), and Monensin (0.5 hours, 200 µM) treatments. Scale bar, 10 µm.

**Figure S11.**
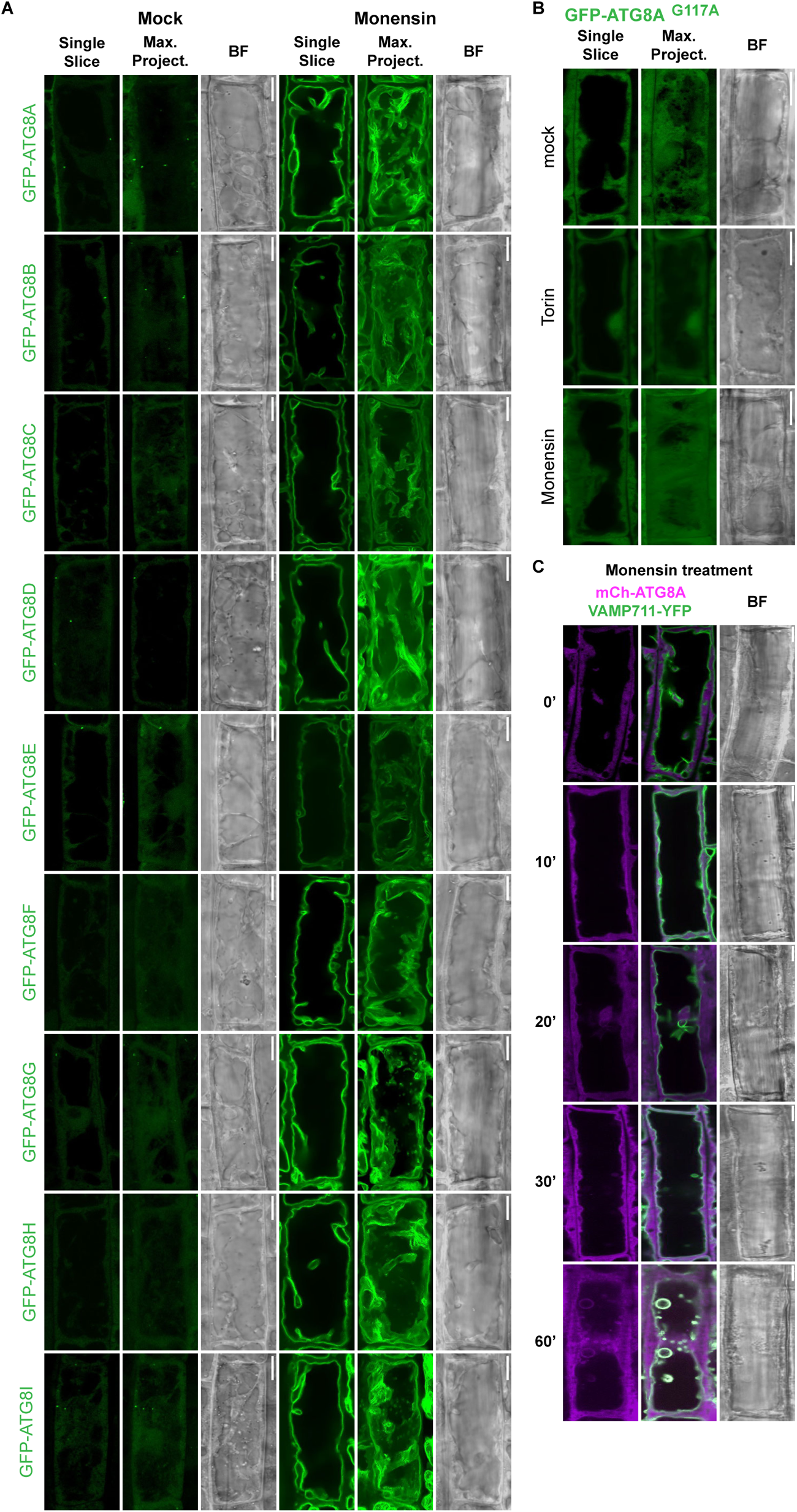
Vacuolar ionophore monensin triggers tonoplast ATG8ylation. **(A)** Confocal micrographs displaying the localization of all nine GFP-tagged ATG8 isoforms (ATG8A to ATG8I) of *Arabidopsis thaliana* under Monensin (0.5 hours, 200 µM) treatment. Scale bar, 10 µm. **(B)** Confocal micrographs of the GFP-ATG8A^G117A^ mutant, highlighting its localization in response to Monensin (0.5 hours, 200 µM) treatment. Images include single optical slices and maximum intensity projections. Scale bar, 10 µm. **(C)** Confocal micrographs of mCherry-ATG8A (in magenta) co-localized with the tonoplast marker VAMP711-YFP illustrating recruitment of ATG8 to the tonoplast upon Monensin treatment. The panel includes a single optical slice alongside a merged image with VAMP711-YFP and a corresponding bright field image. Scale bar, 10 µm. The images follow a time course treatment (0’, 10’, 20’, 30’, 60’) with Monensin (200 µM).

**Figure S12.**
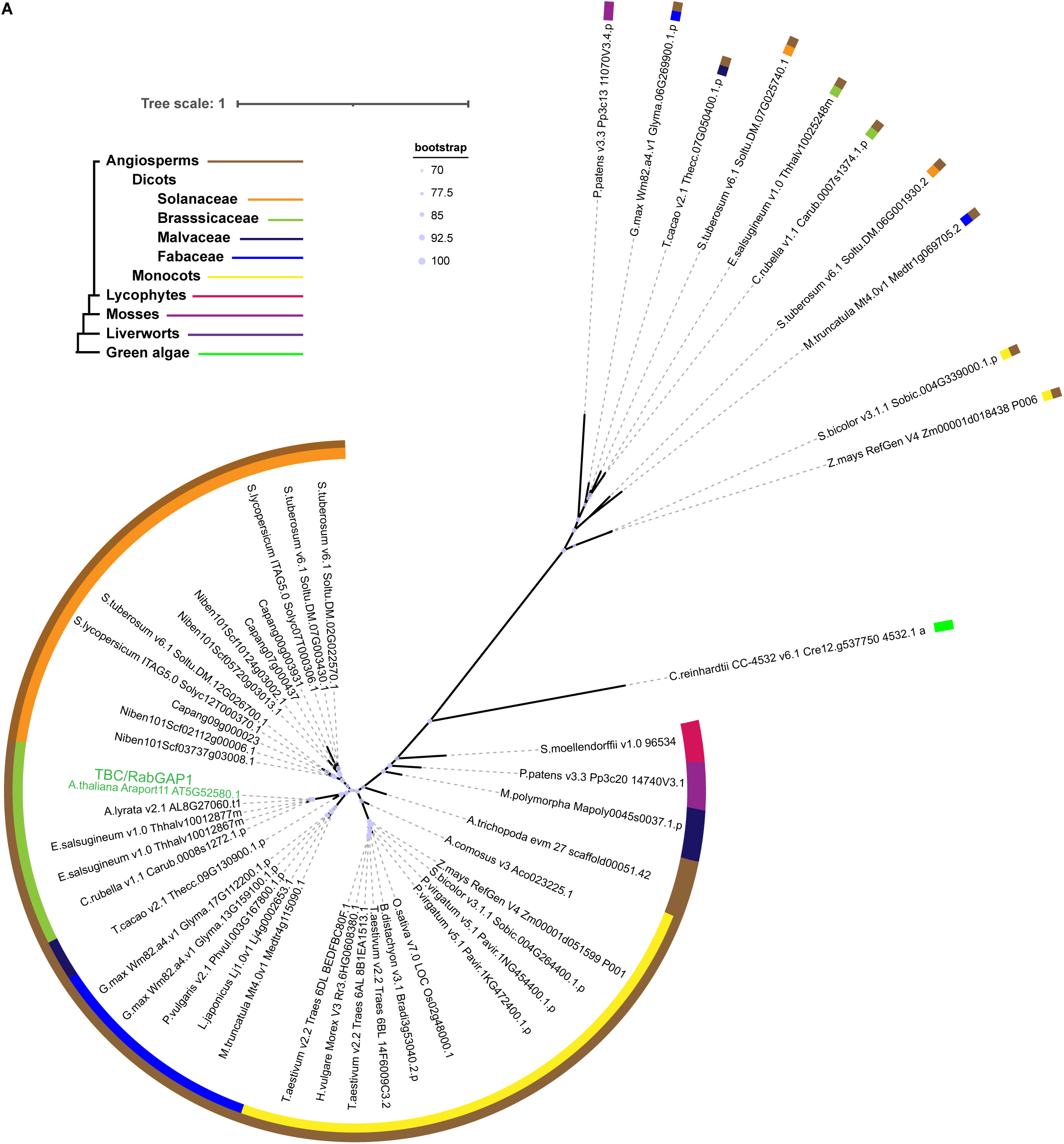
TBC/RabGAP1 is conserved in plants. A plant phylogeny displaying the presence of TBC/RabGAP1 across diverse species. Major plant taxonomic groups are denoted with a colored ribbon. Bootstrap confidence above 70 is shown at the bottom of each new leaf.

**Figure S13.**
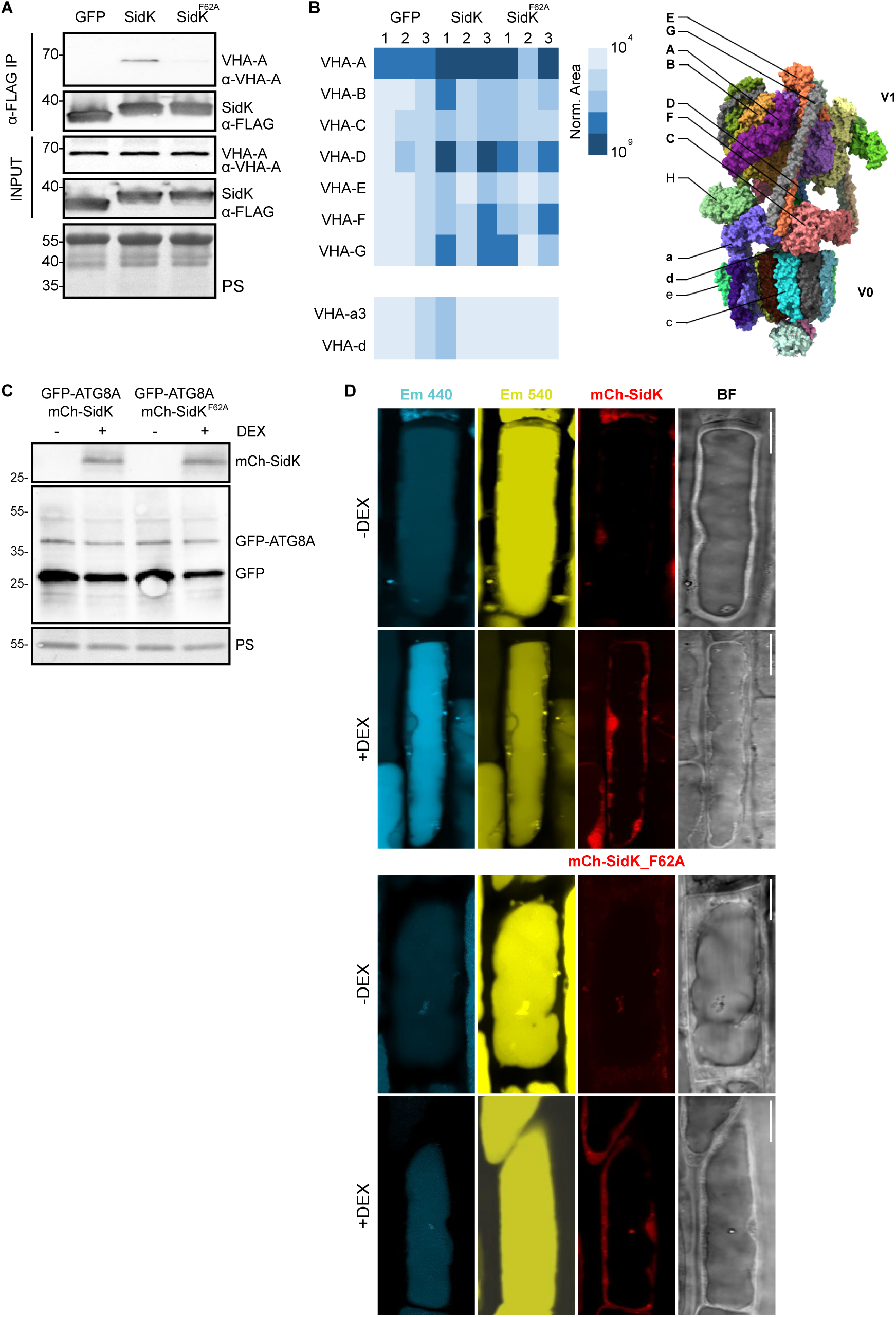
SidK associates with V-ATPase Subunits in *Arabidopsis thaliana*. **(A)** Western blot analysis from immunoprecipitation (IP) experiments using FLAG-GFP, FLAG-SidK, or FLAG-SidK^F62A^ to pull down the VHA-A subunit of the V-ATPase. Blots were probed with anti-FLAG and anti-VHA-A antibodies to detect the presence of VHA-A in the pulldown from each bait protein. INPUTS and anti-FLAG IPs are shown to validate the interaction specificity. **(B)** Immunoprecipitation followed by mass spectrometry (IP-MS) results, presenting the identification of V-ATPase subunits co-immunoprecipitated with FLAG-SidK and FLAG-SidK^F62A^. This panel provides a comprehensive overview of the V-ATPase components pulled down in the experiment detailed in figure S13A, affirming the interaction between SidK variants and the V-ATPase complex. Schematic representation of the V-ATPase complex, detailing all its subunits. The ones marked with bold letters are detected on the SidK IP-MS experiment. **(C)** Inducible expression of SidK as a tool to probe V-ATPase function in Arabidopsis cells. Western blot analysis of GFP-ATG8A lines expressing either mCh-SidK or mCh-SidK^F62A^ mutant under a DEX-inducible promoter, showing protein levels with and without DEX induction. Blots were probed for GFP and mCherry to detect the fusion proteins. **(D)** SidK expression changes vacuolar pH. Confocal micrographs of mCh-SidK or mCh-SidK^F62A^ expressing lines treated with LysoSensor™ Yellow/Blue DND-160, displaying blue, yellow, and red emissions, alongside bright field images, with and without DEX induction. Scale bar, 10 µm.

